# 4R-Tau seeding activity unravels molecular subtypes in patients with Progressive Supranuclear Palsy

**DOI:** 10.1101/2023.09.28.559953

**Authors:** Ivan Martinez-Valbuena, Seojin Lee, Enrique Santamaria, Joaquin Fernandez Irigoyen, Shelley L. Forrest, Jun Li, Hidetomo Tanaka, Blas Couto, Nikolai Gil Reyes, Hania Qamar, Ali M. Karakani, Ain Kim, Konstantin Senkevich, Ekaterina Rogaeva, Susan H. Fox, M. Carmela Tartaglia, Naomi P. Visanji, Tallulah Andrews, Anthony E. Lang, Gabor G. Kovacs

**Author notes:** **Corresponding author:** Gabor G. Kovacs MD PhD FRCPC. University of Toronto, Tanz Centre for Research in Neurodegenerative Disease (CRND), Krembil Discovery Tower, 60 Leonard Ave Toronto On, M5T 0S8, Canada.

## Abstract

Progressive Supranuclear palsy (PSP) is a 4-repeat (4-R) tauopathy. We hypothesized that the molecular diversity of tau could explain the heterogeneity seen in PSP disease progression. To test this hypothesis, we performed an extensive biochemical characterisation of the high molecular weight tau species (HMW-Tau) in 20 different brain regions of 25 PSP patients. We found a correlation between the HMW-Tau species and tau seeding capacity in the primary motor cortex, where we confirmed that an elevated 4R-Tau seeding activity correlates with a shorter disease duration. To identify factors that contribute to these differences, we performed proteomic and spatial transcriptomic analysis that revealed key mechanistic pathways, in particular those involving the immune system, that defined patients demonstrating high and low tau seeding capacity. These observations suggest that differences in the tau seeding activity may contribute to the considerable heterogeneity seen in disease progression of patients suffering from PSP.

## Introduction

Progressive supranuclear palsy (PSP) is a late-onset, four-repeat (4R) tauopathy that belongs to the group of frontotemporal lobar degeneration (FTLD-tau) disorders^1^. Neuropathologically, PSP is characterized by the presence of neuronal loss, neurofibrillary tangles and threads in select subcortical nuclei together with the presence of tufted astrocytes^2–4^. In addition, oligodendroglial coiled bodies and diffuse cytoplasmic immunoreactivity in neurons can be observed^5^. Clinically, the most commonly recognized clinical phenotype of PSP is akinetic-rigid parkinsonism with vertical gaze palsy, axial rigidity, and frequent falls, referred to as Richardson syndrome (PSP-RS)^6^. However, a variety of other clinical presentations of PSP are increasingly recognized^6^.

Tau is a microtubule-associated protein present predominantly in the axons of mature neurons, where it is believed to stabilize microtubules and play a role in axonal transport^7^. Due to its intrinsically disordered nature, tau can adopt a variety of conformations, including β-sheet-rich structures^7^. Converging evidence implies that in tauopathies, a conformational change (misfolding) of the tau monomer produces an aggregation nucleus (seed) that can recruit endogenous tau molecules, and in turn induce their aggregation^7,8^. These tau seeds can self-propagate and progressively spread between inter-connected brain regions exploiting a variety of mechanisms of cell-to-cell transmission^7^.

Intriguingly, a recent study has demonstrated that the initial site of neuronal degeneration and tau pathology in PSP (the “pallido-nigro-luysian” axis and striatum) remains consistent across the different clinical subtypes of the disease^9^. However, it is the differential involvement of other brain regions, where disparities in the amount of tau pathology as well as in the cellular vulnerability patterns (neuronal versus glial), that correlates with the distinct clinical phenotypes^9–14^.

Importantly, how the aggregation of the same protein (tau) can be associated with such variability in the different clinical presentations is not understood. Additionally, differences in the duration of the disease, the clinical presentation, and the presence of diverse concomitant pathologies that present within the same clinical phenotype suggests that there may be potential molecular heterogeneity^15,16^.

In PSP, tau protein accumulates in multiple cell types; however folding appears to be essentially identical from patient-to-patient when evaluated under cryogenic electron microscopy^17^. Recent work in Alzheimer’s disease (AD) has specifically implicated the cytoplasmic soluble, oligomeric species (also known as high-molecular weight tau (HMW-Tau), in the disease pathogenesis^8^. In contrast to fibrillar tau, these smaller oligomers, which are not evident by classical staining or by electron microscopy, can be detected biochemically^8^. These soluble forms of tau are highly heterogeneous in their size distribution and post-translational modification patterns and increasing evidence suggests that such diversity may underlie the heterogeneity seen in neurodegenerative tauopathies^8^. A recent landmark study demonstrated that in AD patient brains, cases which contained more bioactive, propagation-prone material than others were those which presented with the fastest clinical course^18^. The underlying hypothesis of this kind of investigation is that aggregates with a higher seeding ability are likely to have a greater role in the spreading of pathology^19^. One way to identify and study the behavior of these soluble species or seeds is to investigate their seeding capacity *in vitro* using seeding amplification assays (SAAs). These assays capitalize on the property of self-propagation to monitor a seed’s propensity to induce aggregation by amplifying a signal that can sensitively detect minute amounts of these protein seeds using real-time detection of thioflavin T (ThT) fluorescence at multiple time points^20^. We have recently demonstrated the power of this approach to detect differences in the seeding capacity of soluble α-synuclein species isolated from distinct patients with Lewy body disorders (LBD)^21^. In that study, despite similar amounts of α-synuclein pathology in the substantia nigra (SN) of LBD patients with distinct dominant clinical symptoms (motor versus cognitive), the seeding capacity of α-synuclein was dissimilar between LBD subtypes. Interestingly, nigral α-synuclein from patients with rapid disease progression (<3 years) had a higher seeding capacity than the rest of the patients included, in line with what has been described by other groups in AD^18,22^.

Here, to validate the hypothesis that tau soluble species also play a role in PSP clinical heterogeneity, we evaluated 20 different brain regions from 25 PSP patients to detect and quantify these HMW-Tau species. Then, we performed a systematic evaluation of the seeding capacity of patients using 4R-tau SAAs and biosensor cells in one of the brain regions with more HMW-Tau, the primary motor cortex, where striking inter-patient differences in the tau seeding were observed. To further understand the molecular mechanisms leading to these differences in seeding capacity, we performed *in vitro* toxicity assays, neuropathological and biochemical examinations together with a detailed proteotyping and spatial transcriptomics analysis in the primary motor cortex from PSP patients with distinct tau seeding capacities.

## Results

### Mapping and biochemical characterization of HMW-tau in PSP brains

To test the hypothesis that HMW-Tau plays a role in PSP clinical heterogeneity, we mapped the presence of these species in 20 brain regions from 25 subjects with a neuropathological diagnosis of PSP (Supplementary Table 1).

To detect these HMW-Tau species, we immunoblotted under denaturing conditions the proteins present in the PBS-soluble fraction from all the regions and patients using an antibody that recognizes the central region of tau (Tau HT7, Fig. 1a). For quantification purposes, we defined HMW-Tau as the tau with a molecular weight higher than 70 kDa^18^. The values from the quantification of the HMW-Tau (normalized to the expression of β-actin) were obtained from all 25 subjects included and were clustered by brain region (Fig. 1b). Then, they were converted into scores ranging from 0 to 3 representing negative, low, intermediate, or high presence of HMW-Tau and a heatmap was generated to illustrate the diversity of the HMW-Tau species between different brain regions (Fig. 1c).

**Figure 1.**
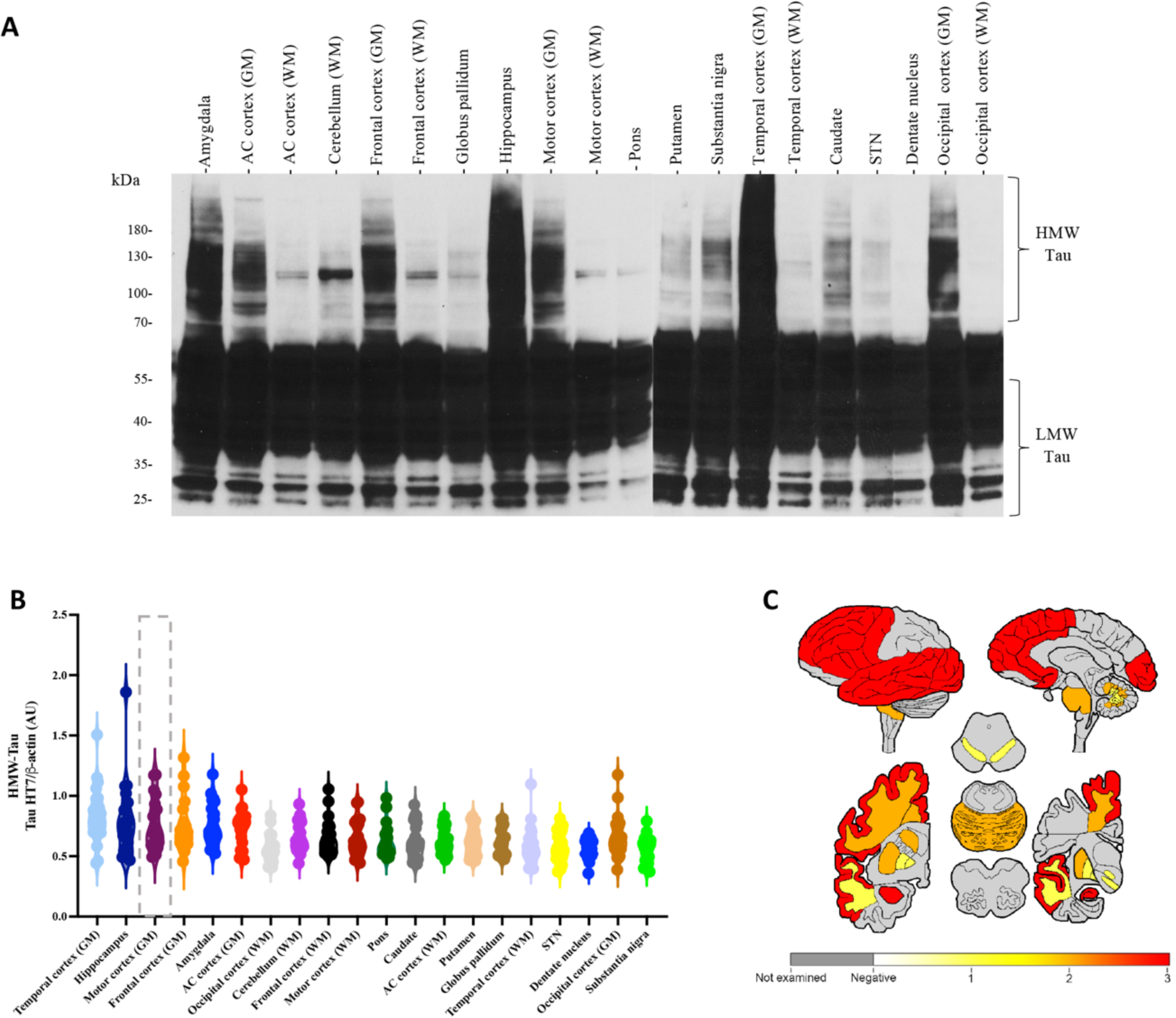
Quantification of HMW-tau reveals extensive heterogeneity across different PSP brain regions. **A)** Representative total tau immunoblots (clone HT7) showing the presence of high (> 70 kDa) and low (< 55 kDa) molecular weight tau from the primary motor cortex PBS-soluble fraction of a PSP patient. **B)** HMW-tau was quantified in all subjects included in the study, clustered by brain region and **C)** converted into scores ranging from 0 to 3 to generate a heatmap. The presence of HMW-tau ranged from yellow (low) to orange (medium) to red (high) in regions evaluated. Grey colored regions indicate that the region was not evaluated. Each dot in (**B**) represents an individual biological sample. HMW: high molecular weight: LMW: low molecular weight. The dashed grey line indicates the brain region that will be used for further analysis.

We found that the temporal cortex and the hippocampus were the two regions expressing the highest quantity of HMW-Tau species, followed by the grey matter of the primary motor (precentral) and middle frontal cortices and the amygdala (Fig.1a-c). Whereas the four regions, which contained the lowest amount of HMW-Tau were the subthalamic nucleus (STN), the cerebellar dentate nucleus, the grey matter of the occipital cortex and the SN. Interestingly, in all five cortical regions examined, HMW-Tau species were always more prominent in the grey matter than in the white matter, with the sole exception of the occipital cortex, where the white matter had more HMW-Tau than the grey matter (Fig. 1a-c). We then correlated the amount of HMW-Tau found in each brain region to the neuropathological PSP stage, we found a significant but negative correlation in the cerebellum white matter (p=0.0382, r=-0.444), in the middle frontal gryus grey (p=0.0095, r=-0.5929) and white (p<0.0001, r=-0.7979) matters and in the putamen (p=0.0056, r=-0.6248).

In addition to the striking differences found within the expression of HMW-Tau amongst the different brain regions examined, we also observed large disparities in HMW-Tau levels among the PSP subjects within the same region (Fig. 1b). Thus, we performed a complete biochemical characterization of the HMW-Tau present in the primary motor cortex. Although the temporal cortex and the hippocampus had higher levels of HMW-Tau, the primary motor cortex is a region heavily involved in the pathophysiology of PSP^9^, and more importantly, is affected by fewer concomitant pathologies, that also involve tau deposition, such as AD or aging-related tau astrogliopathy (Supplementary Table 1).

Thus, we selected 21 PSP cases where frozen primary motor cortex was available and performed a detailed biochemical evaluation of the HMW-Tau. First, we used a panel of three different tau antibodies directed against the N-terminal (Tau 6-18), the central domain (Tau HT7) and the C-terminal (Tau 46) part of the tau protein (Fig. 2a). Interestingly, we observed that the HMW-Tau species were much more evident with the N-terminal tau, in comparison to the amount of HMW-Tau detected using the antibody against the central domain of tau (HT7, Fig. 2a). The C-terminal tau antibody minimally detected these HMW-tau species, but it successfully detected the physiological tau and the low molecular weight tau fragments (Fig. 2a). We then quantified (normalized to expression of β-actin, Fig. 2b) and correlated the HMW-Tau values obtained using our panel of tau antibodies and found a positive and significant correlation between the HMW-Tau species detected with these three antibodies (Fig. 2c).

**Figure 2.**
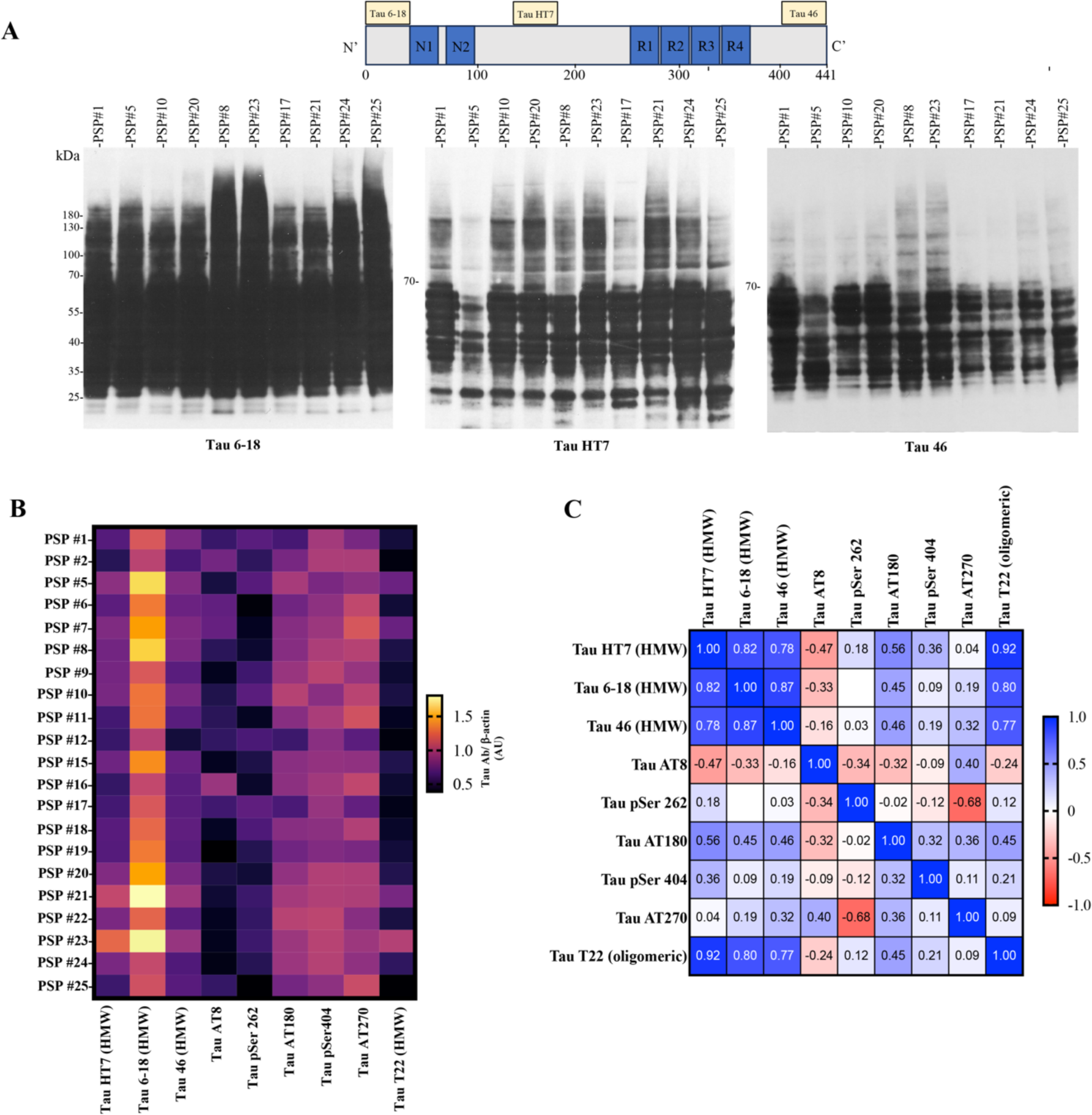
High molecular weight tau is N-terminally enriched, and the tau phosphorylation signature varies across PSP patients. **A)** Representative total tau immunoblots using antibodies against the N terminal (Tau 6-18), the central domain (HT7) and the C terminal (Tau 46) of tau show a distinctive banding pattern of different regions of the tau protein among HMW-tau in the motor cortex of PSP patients. **B)** The HMW-tau assessed with Tau 6-18, HT7 and Tau 46, as well as a panel of antibodies against distinct phospho-tau epitopes and oligomeric tau was quantified and plotted in a heatmap. Yellow colors indicate more expression whereas purple colors indicate less expression. **C)** A matrix is shown to depict the correlations (*r*) between the distinct antibodies used. 1 depicts the strongest correlation, whereas positive or negative values a positive or negative correlation

Next, given the important role that post-translational modifications (PTMs) play in tau aggregation (especially phosphorylation), we used a panel of five widely validated antibodies to evaluate the phosphorylation signature of the tau present in the PBS-soluble fraction. After quantification, we found that tau phosphorylated at threonine 231 (AT270) was the most highly expressed phospho-tau epitope, followed by tau phosphorylated at serine 404 and by tau phosphorylated at threonine 181 (AT180). Interestingly, the two least expressed phospho-tau epitopes in the PBS soluble fraction were tau phosphorylated at serine 202 and 205 (AT8) and at serine 262 (Fig. 2b).

After confirming the inter-individual diversity of tau PTMs between PSP patients (Fig. 2b), we then assessed how the expression of the different phospho-epitopes correlated with the amount of HMW-Tau. We found a positive and significant correlation between AT180 and HMW-Tau (measured using HT7, r=0.56, p=0.008, Fig. 2c). Interestingly AT8 negatively correlated with HMW-Tau (measured using HT7, r=-0.47, p=0.034, Fig. 2c), and tau AT270 negatively correlated with pTau Ser 262 (r=-0.68, p=0.001, Fig. 2c), suggesting a complex interplay between different phospho-epitopes in PSP. Finally, we evaluated whether the HMW-Tau correlated with the amount of oligomeric species found in the PBS-soluble fraction, using a conformational antibody (tau T22^23^). Interestingly, a strong positive correlation was found between the amount of oligomeric tau and HMW-Tau (measured using HT7, r=0.92, p<0.0001, Fig. 2c), suggesting that patients with a high load of HMW-Tau also have a greater presence of tau oligomers.

### 4R-Tau seeding capacity shows great heterogeneity between PSP patients

To evaluate whether the heterogeneity found in the HMW-Tau species impacts tau seeding, we evaluated the seeding capacity from the motor cortex of the 21 PSP patients included in this phase of the study using two complementary approaches: the 4R-Tau SAA and the Tau biosensor cells. 4R-Tau SAAs are cell-free assays that enable the detection of seeding activity from minute amounts of 4R-Tau seeds. To ensure that our assay only recognized 4R-Tau seeds, we first tested it using 1.5μg of total protein derived from the PBS-soluble fraction of the temporal cortex from 2 healthy controls and 2 individuals with AD pathology, together with protein homogenates from the primary motor cortex of 4 PSP patients from our cohort (Supplementary Fig. 2). Our assay detected 4R-Tau seeds in the primary motor cortices of the 4 PSP patients, but no seeding was observed either in the temporal cortex of the controls or in the individuals with AD pathology, validating that the fluorescent signal obtained with this assay was detecting only 4R-Tau seeding.

Having established the specificity of the assay, we evaluated the 4R-Tau seeding capacity from the primary motor cortex. As previously described^21,24^, we quantified 5 kinetic parameters from each reaction: lag time, growth phase, T50 (which corresponds to the time needed to reach 50% of maximum aggregation), ThT max, and area under the curve (AUC) of the fluorescence response. Each kinetic parameter was calculated as the mean of the values obtained from each quadruplicate. We defined the AUC as the seeding parameter of interest as it incorporates all the kinetic features of each aggregation reaction, including the speed and extent of aggregation. The AUC values obtained from the 21 subjects included were clustered (Fig. 3a and Supplementary Table 2) and correlated to the amount of HMW-Tau. There was a robust correlation (r=0.84, p<0.0001) between the presence of higher amounts of HMW-Tau and a higher seeding capacity. However, no significant correlation was found between 4R-tau seeding capacity and age at death (p=0.4768), biological sex (p=0.4765), neuropathological PSP stage (p=0.5915), H1/H2 *MAPT* haplotype (p=0.6358) or *ApoE* genotype (p=0.3579).

**Figure 3.**
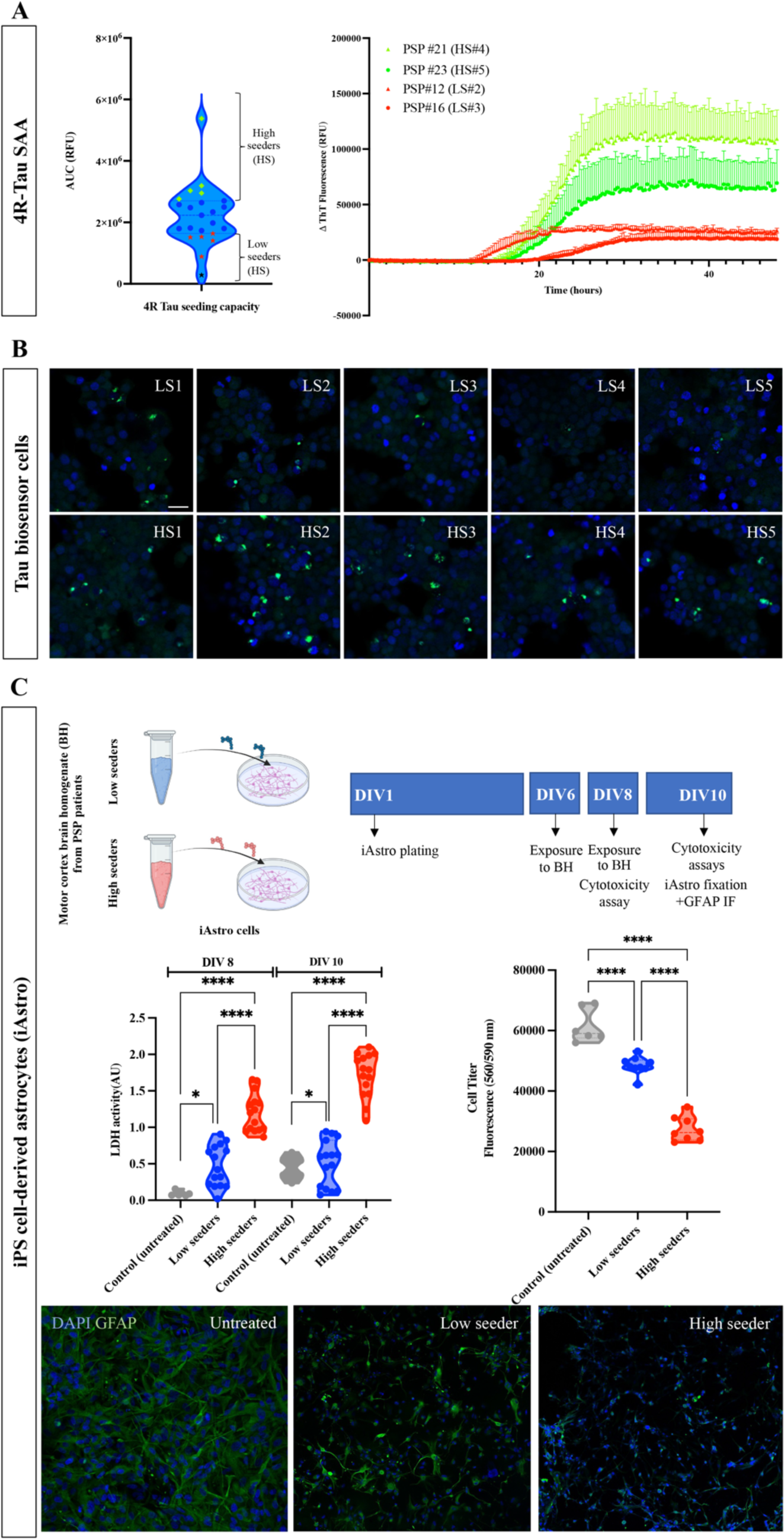
Heterogeneity of tau seeding activity and cytotoxicity in the motor cortex of different PSP patients. **A)** The area under the curve (AUC) from the seeding amplification assay from the primary motor cortex from all the subjects included in the study are shown to display the inter-individual 4R-Tau seeding variability. Each dot represents the mean AUC of an individual biological sample measured in quadruplicate. Green diamonds depict the 5 PSP subjects selected as high seeders and the red asterisks represent the 5 PSP subjects selected as low seeders for further examinations. The blue asterisk depicts the AUC value of PSP#18 that was not included as a part the low seeder group because of its low PSP neuropathological stage. Aggregation curves of 4R-Tau in the presence of motor cortex homogenates from 4 representative PSP patients classified as high and low seeders are also shown. **B)** Representative confocal photomicrographs of tau biosensor cells incubated with the same amount of total protein from the motor cortex brain homogenates derived from high and low seeders showing differential number of inclusions (green inclusions seen as FRET fluorescence). **C)** Schematic of experimental design where human astrocytes derived from induced pluripotent stem cells (iAstro cells) were incubated with PBS-soluble motor-cortex derived homogenates extracted from 10 PSP patients (5 high and 5 low seeders as defined in the 4R-Tau SAA). To evaluate their cytotoxic profiles, at day *in vitro* (DIV) 8 and DIV 10 the concentration of lactate dehydrogenase (LDH) was measured. In addition, at DIV10 a cell viability assay (CellTiter-Blue®) was performed to estimate the number of viable cells. Representative confocal photomicrographs of iAstro cells incubated with primary motor cortex brain homogenates derived from high and low seeders and immunostained with a GFAP antibody were acquired to complement the cytotoxic assays.

We next subtyped the cases into 3 groups: high seeders (those whose AUC values fell within the 3^rd^ quartile), intermediate seeders, and low seeders (those whose AUC values fell within the 1^st^ quartile), according to their ability to misfold monomers of recombinant Tau *in vitro* (Fig. 3a). To assess whether these distinctions could be due to differences in seed concentration or seed characteristics, we performed end-point dilution of the primary motor cortex PBS-soluble fraction from two PSP cases, one classified as a high seeder and the other a low seeder (Supplementary Fig. 3). Regardless of the amount of brain homogenate used to seed the 4R-Tau SAA reaction, the seeding activity of the misfolded tau from the high-seeder PSP case was more potent than that from the low-seeder PSP case. Although we did not find any significant differences between high and low seeders with regards to the PSP neuropathological stage, PSP clinical phenotype, concomitant co-pathologies, sex, *MAPT* haplotype or *ApoE* genotype, the age at death and disease durations were different. The high seeders, on average, were younger than the low seeders (71.6 versus 76.8 years), and, also had a shorter duration of disease (4.4 years) compared to the low seeders (9.2 years, Supplementary Table 1). To illustrate the different profile of 4R-Tau SAA of high and low seeders we plotted SAA data from the primary motor cortex of two representative low and two high seeder cases (Fig. 3b).

To validate our 4R-Tau SAA results, we measured tau seeding bioactivity using a widely used Forster resonance energy transfer (FRET) based biosensor cell line^18,25^. We exposed the cells to 6μg of total protein derived from the PBS-soluble fraction of the primary motor cortex of the same 21 PSP cases for 48 hours. After the incubation period, we quantified the percentage of FRET positive inclusions in these cells using an image-based analysis (Supplementary Fig. 4 and Supplementary table 2). Interestingly, the number of positive FRET inclusions significantly correlated with the 4R-SAA AUC values (r=0.485, p=0.0243), confirming that there is heterogeneity of tau seeding between PSP patients, specially between those categorized as high and low seeders (Fig. 3b).

To evaluate whether the primary motor cortex homogenates derived from high and low seeders also have different cytotoxic profiles that could explain the discrepancy in the duration of the disease, we used commercially available human astrocytes derived from induced pluripotent stem cells (iAstrocyte). To match the amount of protein added to the tau biosensor cells, the iAstrocytes were exposed to 6μg of total protein from the PBS-soluble fraction of the primary motor cortex for 4 days (Fig. 3c). To evaluate the cytotoxic effect of primary motor cortex-derived homogenates, we used two different approaches^21^. First, at day *in vitro* (DIV) 8 and DIV 10 we measured the concentration of lactate dehydrogenase (LDH), as damage of the plasma membrane results in a release of this cytosolic enzyme into the surrounding cell culture medium. When we measured extracellular LDH, we found that at both DIV8 and DIV10 LDH was higher in the cells treated with motor cortex homogenates derived from either high or low seeder patients compared to untreated cells (p<0.0001, Fig. 3c). However, when we compared the LDH levels between PSP patients, we found significantly higher (p<0.0001) extracellular LDH expression from the cells exposed to the primary motor cortex homogenates derived from the high seeder patients compared to low seeder patients (Fig. 3c). To corroborate these findings, we performed a cell viability assay (CellTiter-Blue®) at DIV10, where we found a significant decrease in cell viability in iAstrocytes cells exposed to primary motor cortex homogenates derived from high seeders, compared to those exposed to primary motor cortex homogenates derived from low seeders (p<0.0001, Fig. 3c). At DIV 10, neurons were fixed and immunostained using the astrocytic maker glial fibrillary acidic protein (GFAP, Fig. 3c). We observed qualitatively that the iAstrocytes cells exposed to primary motor cortex homogenates derived from low seeders were fewer compared to untreated astrocytes (Fig. 3c). Furthermore, iAstrocytes in the wells treated with primary motor cortex homogenates derived from high seeder patients were scarce (Fig. 3c). These results extend the findings obtained with the 4R-Tau SAA and suggest that the tau and other co-factor/proteins present in PSP patients possess not only a different seeding capacity, but also confer different cytotoxic properties.

We further characterized tau in the PBS-soluble homogenates from the primary motor cortex of 10 patients (5 classified as high and 5 as low seeders) using two protease-sensitivity digestion assays that had been previously used to define conformations of amyloidogenic proteins^18,26^. Primary motor-cortex homogenates were incubated with proteinase K (PK, 20μg/ml) and thermolysin (TL, 25μg/ml). Although the protein extracts from patients with PSP had similar tau banding patterns when immunoblotted under denaturing conditions (Fig. 2a), these banding patterns markedly differed between the high and the low seeders after treatment with PK and TL (Fig. 4). The most divergent PK and TL pattern was observed with the tau antibody against the N-terminal domain (Tau 6-18, Fig. 4a). Three out of the five high seeder cases displayed numerous PK and TL resistant bands that were not evident in the low seeder cases. Furthermore, a more prominent 27kDa band was observed in the high *vs* low seeders. This latter finding was also consistent when the PK and TL digested samples were probed against two other tau antibodies (Tau HT7 and Tau46) (Fig. 4b and c). These results indicate both striking inter-individual differences in protease sensitivity between subjects with PSP and that the tau derived from the high seeder patients is more resistant to PK and TL digestion.

**Figure 4.**
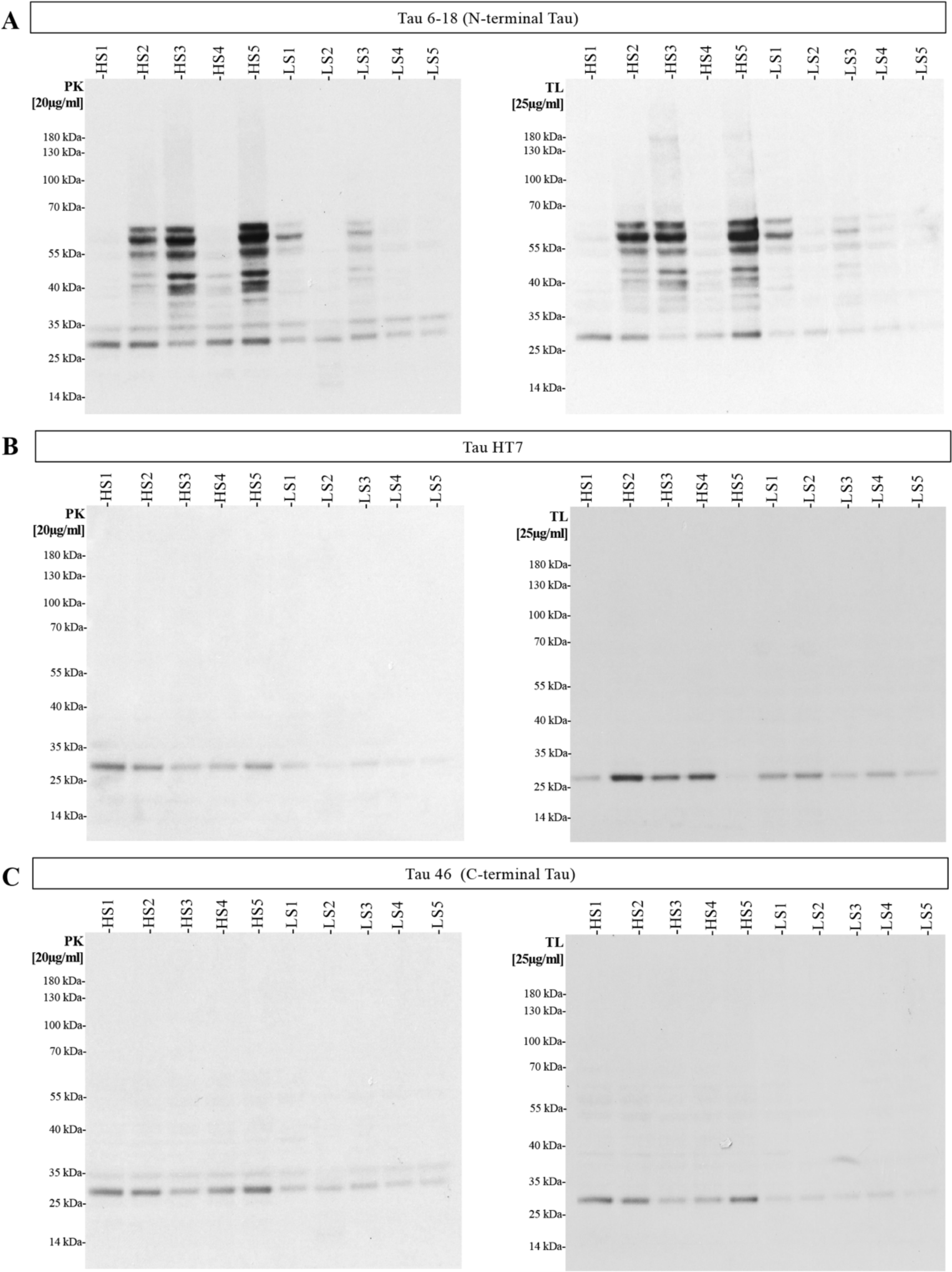
Tau derived from high and low seeders PSP cases has distinct protease resistance profiles. Representative immunoblots of primary motor cortex samples subjected to either proteinase K (PK) or thermolysin (TL) digestion show different profiles of **A)** N-terminal **B)** central domain and **C)** C-terminal tau regions.

### Proteotype remodeling between PSP cases with different 4R-Tau seeding capacity

To elucidate whether the diversity in HMW-Tau species and 4R-Tau seeding capacity could have an impact on the molecular and cellular mechanisms that might explain the differences in the duration of disease between PSP patients, we performed mass-spectrometry-based quantitative proteomics. We analyzed the primary motor cortex proteome of 9 PSP patients (4 high seeders and 5 low seeders) which were used for the *in vitro* experiments (due to tissue unavailability HS#2 was excluded from the proteomic analysis). We identified 3,767 proteins in the primary motor cortex of these patients, of which 484 were differentially expressed in high *vs* low seeders (Fig. 5a); 247 were more highly expressed and 237 had lower expression in the high vs low seeders. Plotting the differentially expressed proteins (DEPs) demonstrated one cluster involving the four high-seeder cases, another very separate cluster involving three of the low-seeder cases (LS 1, 4 and 5), and a third different proteomic signature in the other 2 low seeders (LS 2 and 3) (Fig. 5a). Despite the differences in the proteomic signature between the 5 low seeder cases, we decided to perform the subsequent bioinformatic analysis with the 4 high and the 5 low seeder cases.

**Figure 5.**
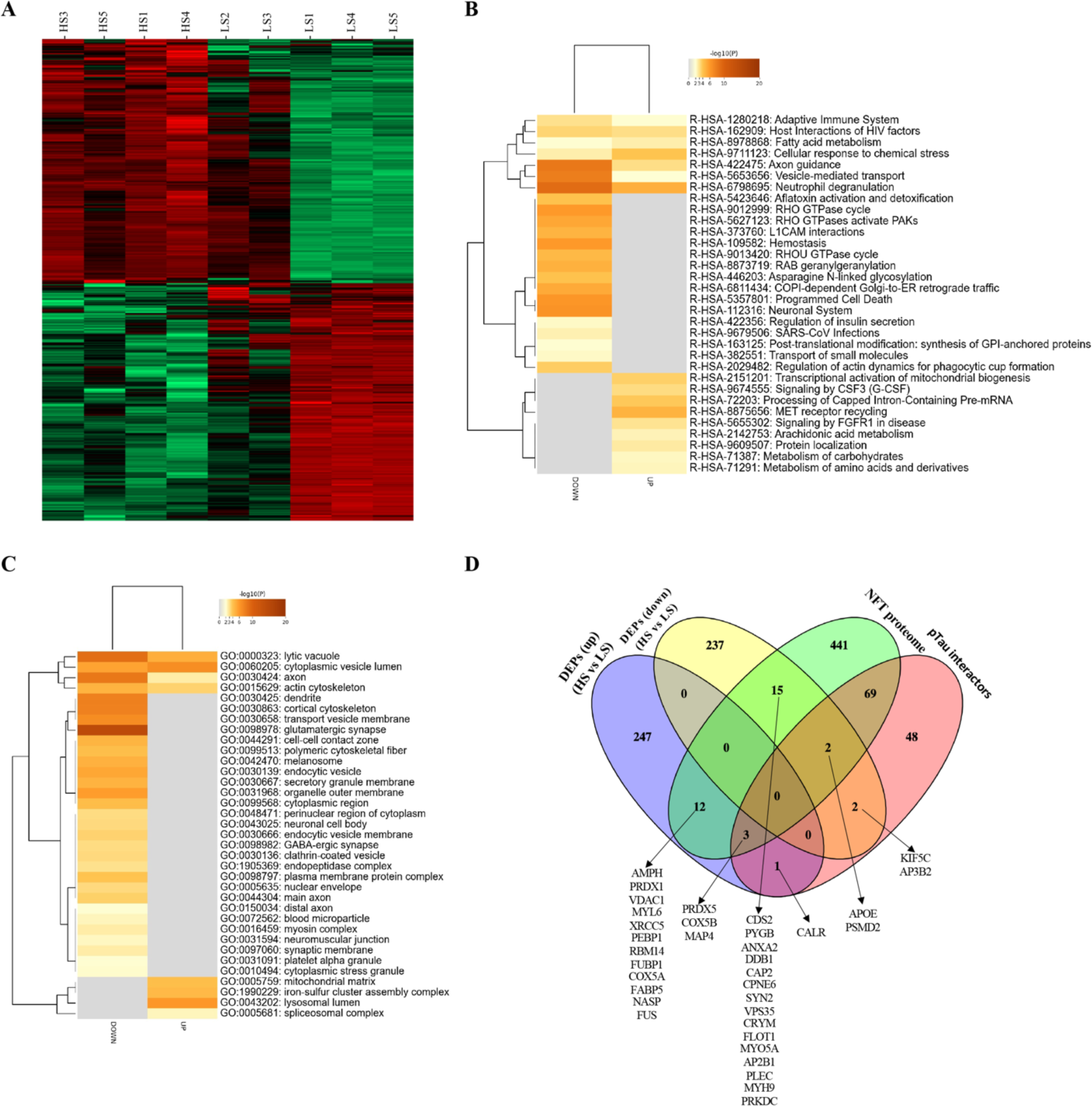
Primary motor cortex proteome remodeling in PSP cases with high and low tau seeding activity. **A)** Heatmap showing both unsupervised clustering and the degree of change for the differentially expressed proteins (DEPs). **B)** DEPs were independently analyzed to map common pathways implicated by deregulated proteins as well as specificities in each differential protein subset **C)** DEPs were also independently analyzed to map altered proteins in specific subcellular compartments. **D**) A Venn diagram highlighting the overlap between the DEPs and known phospho-tau interactors, as well as proteins identified in neurofibrillary tangles.

We examined the differential proteome distributions across specific pathways/biofunctions (Fig. 5b, Supplementary Fig. 5). Up- and down-regulated proteins were independently analyzed. Proteins related to the adaptative immune system, as well as to the metabolism of fatty acids, axon guidance and to the cellular response to chemical stress were both up- and down-regulated when comparing high and low seeder patients. However, proteins linked to hemostasis, programmed cell death, regulation of insulin secretion and transport of small molecules were significantly downregulated in high *vs* low seeders. In contrast, proteins related to the transcriptional activation of mitochondrial biogenesis, protein localization, or to the metabolism of carbohydrates and amino acids were significantly upregulated in the high seeders (Fig. 5b). We also performed a subcellular mapping of the DEPs. As for the pathway analysis, up- and down-regulated proteins were independently analyzed. We found that up- and down-regulated proteins had an impact on different cellular structures, and whereas in lytic vacuoles, axons and actin cytoskeleton, both up- and down-regulated proteins were found, upregulated proteins were only found in four subcellular compartments: the mitochondrial matrix, the lysosomal lumen, the iron-sulfur cluster assembly complex and the spliceosomal complex. However, down-regulated proteins were found in many subcellular compartments, including dendrites, endocytic vesicles, neuronal cell body, the nuclear envelope and in the main and distal axons (Fig. 5c).

Finally, we performed a network analysis to evaluate whether any of the DEPs had been previously associated with tau in the literature. Our analysis revealed that of the 484 dysregulated proteins, 8 proteins had been reported to functionally interact with tau (Fig. 5d). Interestingly, our analysis also revelated that 27 of our DEPs had also been found in the phosphorylated tau interactome of AD neurofibrillary tangles^27^ (Fig. 5d).

### Cases with different 4R-Tau seeding capacity can be neuropathologically clustered

Based on the distinct proteomic profile and the marked differences observed in the 4R-Tau seeding capacity, we explored the possibility of demonstrating the presence of distinctive neuropathological features by assessing two high seeder (HS 3 and 5) and two low seeder (LS4 and 5) cases. These four cases were not different in the stage of PSP (stage four/five), clinical phenotype (PSP-RS) or in the presence of co-pathologies (Supplementary Table 1). The neuropathological examination was performed by two neuropathologists (S.L.F and G.G.K) blinded to the seeding category of the cases. A thorough evaluation of neuronal, astroglial and oligodendroglial tau-immunopositive inclusions throughout the brain of the four cases revealed a similar distribution and density of total tau-immunopositive pathology (Fig. 6a). However, the density of each cytopathology differed between regions and cases. In all cases, the density of neurofibrillary tangles was similar, with the highest density observed in the brainstem and globus pallidus (Fig. 6a). In contrast, heat mapping showed variation in the distribution and density of tufted astrocytes and oligodendroglial coiled bodies (Fig. 6b) both between individual cases, and between high *vs* low seeders. Overall, the highest density of tufted astrocytes was observed in the primary motor cortex, basal ganglia and thalamus, and the highest density of oligodendroglial coiled bodies was observed in the primary motor cortex, basal ganglia, thalamus, brainstem and cerebellar white matter. A high density of tufted astrocytes was also found in the parietal cortex of high seeders. Although a clear differentiation between low and high seeders could not be determined when visualising the neuropathological features, cluster analysis based on the semi-quantitative scores for all brain regions and all cytopathologies (neuronal, astroglial, oligodendroglial) revealed that the two low seeders cluster together and have a distinct pathological profile compared to the high seeders (Fig. 6c).

**Figure 6.**
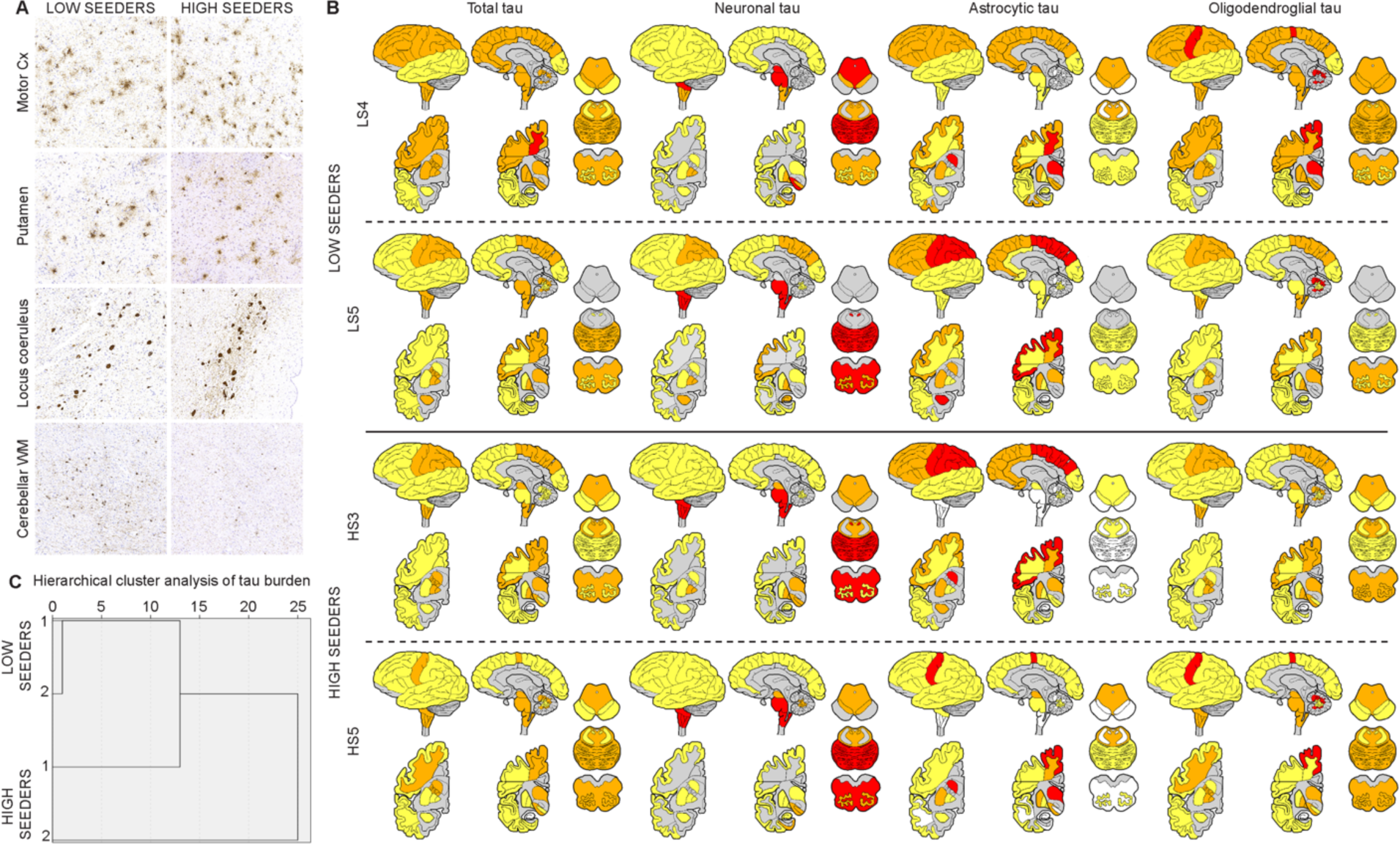
Comparison and distribution of tau-immunopositive pathology in major brain regions of low and high seeders. **A)** Images of LS#5 are on the left and images of HS#5 are on the right panel. Neurofibrillary tangles, tufted astrocytes and coiled bodies in the primary motor cortex and putamen. Neurofibrillary tangles in the locus coeruleus and oligodendroglial coiled bodies in the cerebellar white matter (WM). **B)** Heat map images of total tau, neuronal, astroglial and oligodendroglial tau pathology in the two low and high seeders. Each case examined are represented horizontally, with the two low seeders (LS#4 and 5) at the top and the two high seeders (HS#3 and 5) represented at the bottom. Total tau, neuronal, astroglial and oligodendroglial tau-immunopositive pathology are represented in each vertical column. Red = high, orange = moderate, and yellow = low density of tau pathology. White = absence of pathology. Grey = areas not examined. **C)** Cluster analysis based on all tau-immunopositive cytopathologies across brain regions reveal that the two low seeders cluster together and have a distinct pathological profile to the high seeders.

### Spatial transcriptomics confirms molecular differences between PSP subgroups

After confirming that besides the differences found in 4R-Tau seeding and in the proteomic signature, the high and low seeders could be neuropathologically clustered based on their seeding capacity, we performed spatial transcriptomic analysis in the same four cases where a thorough neuropathological examination had been performed. After confirming that the four cases had good tissue integrity and RNA quality (Fig. 7a), transcriptomic data was integrated and clustered using standard pipelines, revealing 11 distinct region-types within the motor cortex. All clusters expressed neuronal markers, with highest expression in Clusters 2-5 (Fig. 7e). Clusters 0,1, and 6 additionally expressed oligodendrocyte markers. Cluster 7 additionally expressed endothelial and microglia markers. Cluster 8 had the highest expression of astrocyte markers.

**Figure 7:**
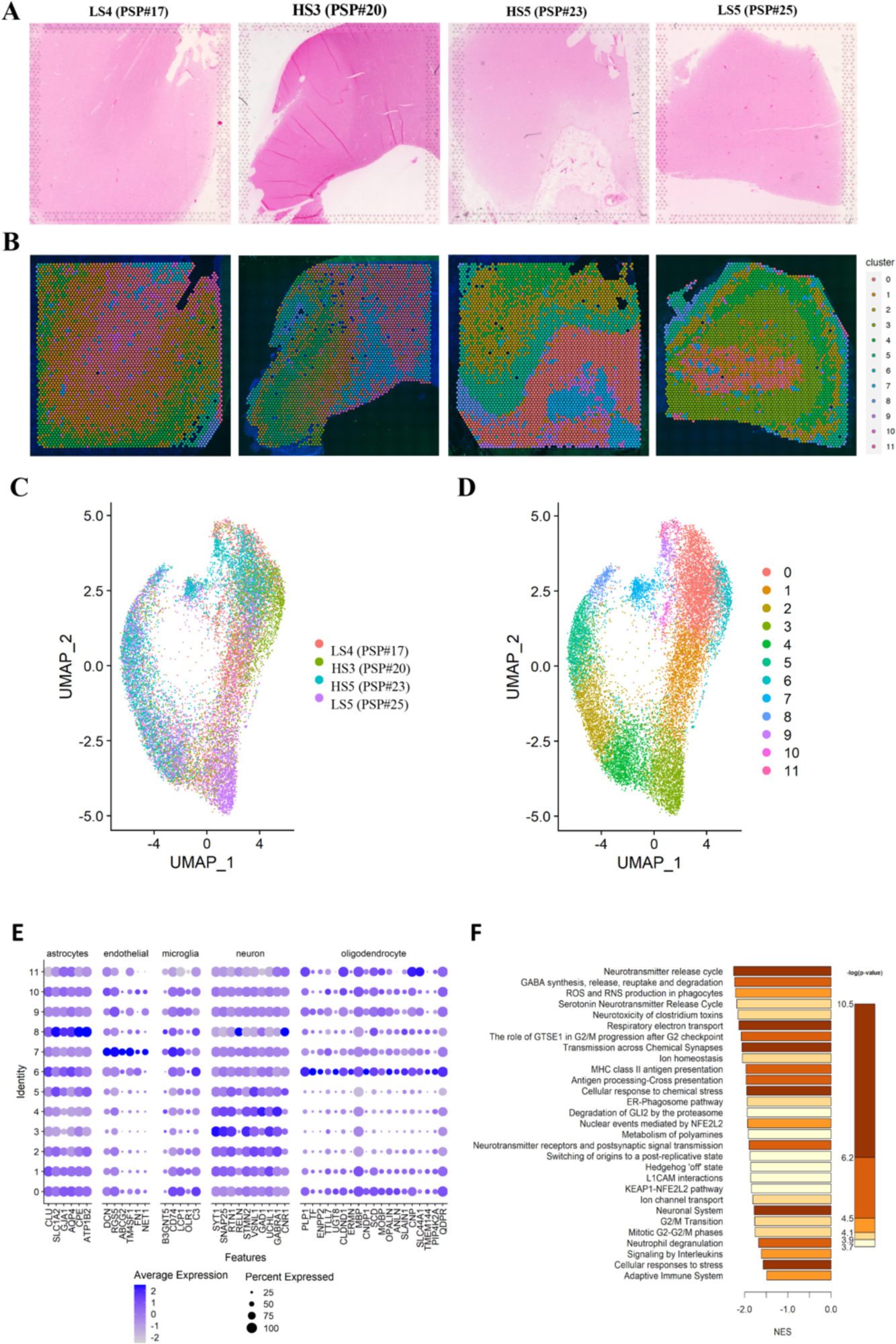
Spatial Transcriptomics confirms immune and neuronal dysregulation in high-seeder PSP. **A)** Two high-seeder and two low-seeder PSP samples were stained with H&E and subject to VISIUM spatial transcriptomics. **B)** Clustering of integrated data revealed 11 distinct spatial domains across all four slices. **C)** Uniform Manifold AProximation (UMAP) confirms integration of spatial transcriptomic profiles across slices, and **D)** presence of 11 distinct clusters. **E)** Expression of cell-type marker genes reveal compositional differences between spatial domains. **F)** Top 30 significantly enriched Realtime pathways among differentially expressed genes overlap with those detected using proteomics.

Each tissue slice contained all regions but in differing proportions and arrangements (Fig. 7b-d) due to slicing orientations and locations within the motor cortex. Thus, we did not find any significant differences in the frequencies of these clusters between high- and low-seeders.

We then performed differential expression analysis between high and low seeders controlling for the proportions of different spatial clusters (2 samples per group). Pathway enrichment analysis across all genes revealed many enriched pathways downregulated in high *vs* low seeders at the transcriptomic level that are consistent with those observed at the protein level, including the Adaptive Immune System, L1CAM interactions, Neuronal System, and Neutrophil degranulation (Fig. 7f). Transcriptomic analysis further revealed downregulation of the cell-cycle, and disruption of serotonin and GABA release pathways in the high-seeder PSP cases.

### Misfolded tau accumulation produces transcriptomic changes

Finally, and taking advantage of the spatial resolution that Visium offers, we next examined whether these transcriptomic changes were associated with the aggregation of tau. Using an innovative approach for the use of spatial transcriptomic studies in diseases involving protein deposition, AT8-tau load was measured in an adjacent tissue section, quantified using Halo, and aligned with the transcriptomic data (Fig. 8a). Using a general linear model, we identified gene expression patterns associated with varying concentrations of tau across 1,250 Visium regions of interests (ROIs) from one high-seeder and one low-seeder sample which exhibited fine alignment with the IHC slide. This identified 69 genes significantly associated with tau misfolding, where 58 were up-regulated in proximity to tau and 11 were down-regulated (Fig. 8b and Supplementary Table 3). In agreement with our differential expression between low and high seeders, we found these genes were enriched in negative regulation of the cell-cycle, negative regulation of transcription factor binding, and extracellular matrix organization (Fig. 8c).

**Figure 8.**
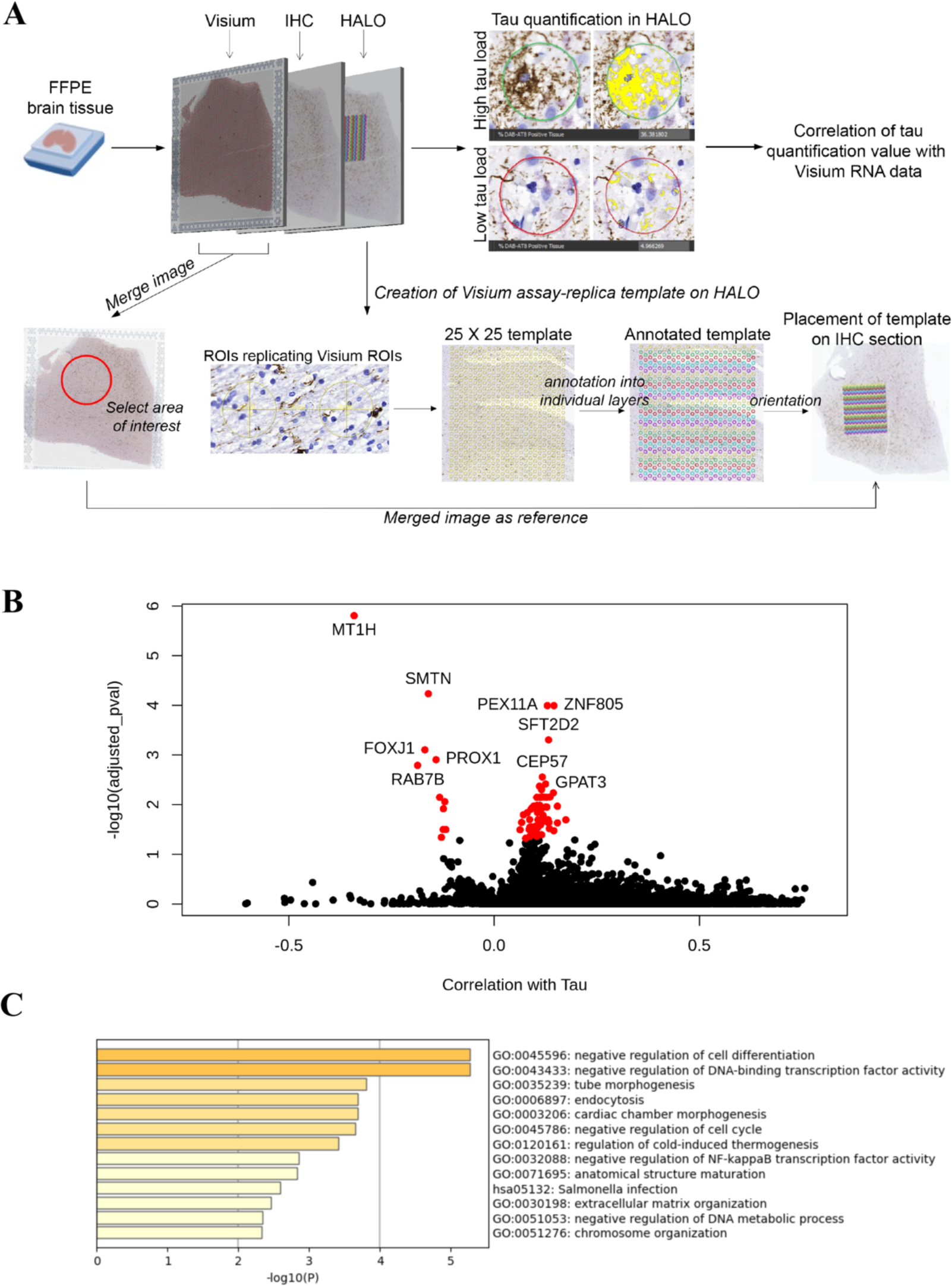
Dysregulation of cell-cycle and stemness pathways is associated with tau protein aggregates. **A)** Adjacent tissue slices were used to quantify tau expression and align quantification estimates with VISIUM RNA capture spots using Halo. **B)** 69 genes were significantly differentially expressed in proximity to misfolded tau aggregates. **C)** Gene Ontology (GO) functional enrichments reveal dysregulation of cell-cycle and stemness pathways associated with tau protein aggregation

## Discussion

Our study has revealed several novel aspects of PSP pathophysiology linked to the differential 4R-Tau seeding capacity found in a cohort of clinically and neuropathologically deeply phenotyped cases. First, we have provided evidence of a previously unrecognized heterogeneity in the presence of HMW-Tau species across 20 different brain regions. Second, we have linked the varying expression of these species among patients to the amount of tau oligomers and to a differential 4R-Tau seeding capacity. Third, we have confirmed that an elevated 4R-Tau seeding activity correlates with both higher cytotoxicity *in vitro* and corresponds with a shorter disease duration. Fourth, the differentiation of cases based on the 4R-Tau seeding capacity aligns with clustering of PSP cases based on the constellation of neuropathological variables across brain regions irrespective of PSP neuropathological stage. Finally, our proteomic and spatial transcriptomic analysis have identified a significant disruption in the adaptative immune system in high *vs* low seeder patients. For the first time, we have also demonstrated a correlation between tau deposition and a negative regulation of the cell-cycle, as well as a negative regulation of transcription factor binding and the extracellular matrix organization.

PSP is a clinically heterogeneous tauopathy, and although the regional distribution of 4R-Tau pathology might correlate with the predominant clinical features^9–14^, the burden of tau inclusions does not fully explain the significant differences in clinical presentation and rate of disease progression exhibited in this disease^16^. Accumulating evidence in other neurodegenerative diseases, like LBD, multiple system atrophy (MSA) and AD, indicates that the conformational diversity of α-synuclein and tau might explain part of the clinical heterogeneity found in these diseases^18,21,24^. Much of the recent work in the tau field has focused on elucidating which form of tau is the first to get misfolded^28,29^ and which form(s) of tau are triggering the propagation of the pathology across the brain^30^. In AD, a handful of studies have implicated cytoplasmic soluble, oligomeric species, that are highly heterogeneous in their size distribution and PTM patterns, in the clinical heterogeneity seen in this disease^8^. Thus, our first objective was to evaluate whether these cytoplasmic soluble species were also present in the brain of PSP patients. Using western blotting, we performed the first comprehensive mapping of HMW-Tau in the PBS-soluble fraction of 20 different brain regions from 25 neuropathologically-confirmed PSP cases. We found that these soluble species were present in all regions examined, although with a high degree of variability between different brain regions and patients. The temporal cortex and the hippocampus were the regions where more of these species were found, probably due to the presence of concomitant AD pathology in these brain regions. However, our analysis revealed that early affected regions in PSP^9^, such as the STN, the SN or the globus pallidus did not have a high presence of HMW-Tau. An explanation for this interesting finding could be that over the course of the disease, these soluble tau species, present at the earlier stages of the disease, would become sequestered in larger insoluble aggregates at later stages of the disease^31^.

Besides the high diversity of HMW-Tau levels between regions, our brain mapping also showed great heterogeneity among patients. Using the primary motor cortex as region of reference for our analysis, we conducted a biochemical characterization of these HMW-Tau species. We first evaluated whether these species could be detected using antibodies directed against the C- and the N-terminal domains of tau. Interestingly, we found that while the antibody directed against the N-terminal domain of tau was able to detect more HMW-Tau than the one used for the HMW-Tau brain mapping (directed against the central domain of tau), and when the C-terminal antibody was used, the amount of HMW-Tau species detected was negligible. These results could suggest that the fold of the HMW-Tau species might expose more of the N-terminal and the central domains of tau than the C-terminal part, although further structural analysis is needed to validate these findings.

Due to the important role of PTM in tau misfolding^32,33^, we next assessed the tau phosphorylation signature of the PSP patients included in our study. Interestingly, we found that amongst the five epitopes probed in our panel of phospho-tau antibodies, those which detect the phosphorylation of tau at serine 202 and 205 (AT8) showed the least expression in the PBS soluble fraction. However, AT8 was not only the least expressed among the phospho-tau antibodies evaluated, but it also negatively correlated with the levels of HMW-Tau. AT8 is one of the most widely used antibodies to detect tau pathology in neuropathological practice^9,34^ because it labels all types of 4R-Tau related pathology^5,9^, which might indicate that in PSP, the tau phosphorylation in its serine 202 and 205 is found in more advanced stages of tau aggregation. Although this result needs further validation in larger cohorts of PSP patients and the role of serine 202 and 205 phosphorylation in tau misfolding and aggregation needs to be further explored, this finding is in line with previous work from our group, where the amount of aggregated α-synuclein (detected using the 5G4 antibody) negatively correlated with the α-synuclein seeding capacity in LBD^21^ and MSA patients^24^. To evaluate whether the higher presence of HMW-Tau had any correlation with tau seeding efficacy, we evaluated the Tau seeding capacity using two methods: 4R-Tau SAA and tau biosensor cells. The capacity to monitor in real time how tau aggregates template the misfolding of recombinant tau fragments has been previously exploited to generate SAAs that are able to measure 4R-Tau seeding kinetics in different biological samples^35^. Using this assay, we analyzed the primary motor cortex 4R-Tau seeding activity in all patients included in the study. The results obtained using the 4R-Tau SAA were further validated using a complementary and widely validated method that also allows the evaluation of tau seeding capacity in cultured biosensor cells^25^. Following these assays, we subclassified the PSP cases into 3 groups: high, intermediate and low seeders, according to their ability to misfold recombinant 4R-Tau fragments and following the same classification that has previously been applied in AD^18^, MSA^24^, and LBD^21^ patients. To further elucidate the underlying mechanisms behind the differential tau seeding activity seen in PSP patients, we selected 5 high seeders and 5 low seeders. In these 10 PSP patients we performed *in vitro* cytotoxicity assays followed by a proteomic examination of the primary motor cortex. For the cytotoxicity assays, we used human astrocytes derived from induced pluripotent stem cells. After we incubated these cells with brain extracts derived from the primary motor cortex of these 10 PSP patients, we found a striking decrease in cell viability in the astrocytes exposed to high seeder-derived primary motor cortex homogenates compared to those exposed to the homogenates derived from low seeders. Although these findings might explain the finding that a high 4R-Tau seeding capacity correlates with a rapid disease progression, further studies will be required to validate this key finding.

One unavoidable limitation of our work is that the extracts from PSP brains also contained other proteins, in addition to the pathological tau, that could be responsible for the differential cell death rates. Thus, we used two different protease-sensitivity digestion assays that differentiate protein conformations^18,26^, to evaluate whether any structural differences in the HMW-Tau could be found between high and low seeders. Interestingly, we found that the tau from high seeder PSP patients was more resistant to PK and TL digestion than the tau found in the low seeders. Furthermore, we also found different PK and TL banding patterns between PSP subjects when the digested tau fragments were probed against an N-terminal tau antibody, further strengthening the necessity of structural analysis of these species. Altogether, these data support the hypothesis that different biologically active conformers of tau exist across PSP brains, raising the intriguing possibility that different underlying pathophysiological mechanisms might occur in different subsets of patients, representing a biological basis for differences in the clinical course.

To test this hypothesis, we performed mass-spectrometry-based quantitative proteomics and analyzed the primary motor cortex proteome of 9 PSP patients that were previously used for the *in vitro* experiments. Our analysis found that 484 proteins were differentially expressed in high *vs* low seeders. Interestingly, we found that lytic vacuoles, axons, the lysosomal lumen or the mitochondrial matrix were the most affected subcellular compartments. Furthermore, an analysis to explore the differential proteome distributions across specific pathways/biofunctions revealed that most DEPs were related to changes in the adaptive immune system, the cellular response to chemical stress, programmed cell death or mitochondrial biogenesis. Our analysis also revealed that from the 484 dysregulated proteins, 8 proteins (including APOE^36^) had been reported to functionally interact with tau and 27 proteins (including FUS^37^) had also been found in the phosphorylated tau interactome of AD neurofibrillary tangles^27^.

However, it is important to note that despite highlighting processes consistent with current concepts of neurodegenerative disease, a major limitation of the current approach is that we are unable to determine whether the observed changes in the proteome are related to the tau deposition. When using many -omics technologies in bulk tissue homogenates, there is a loss of spatial information, including the relationship of cells to tau deposition^38,39^. To overcome this limitation, we applied a novel technology that uses spatially barcoded arrays, allowing unbiased transcriptome profiling in tissue sections, maintaining the spatial localization of the sequenced molecules^38^. This spatial transcriptomic analysis in the primary motor cortex of four patients (two high and two low seeders) also showed a very distinctive proteomic signature.

Clinically, these cases were all PSP-RS and had a similar PSP neuropathological stage. Notably, two neuropathologists performed a blinded neuropathological examination of these four cases, and when a cluster analysis based on the semi-quantitative scores for all brain regions and all cytopathologies (neuronal, astroglial, oligodendroglial) was performed, it revealed that the two low seeders clustered together and had a distinct pathological profile compared to the high seeders.

Interestingly, differential expression analysis and pathway enrichment analysis, showed many enriched pathways up- and down-regulated in high seeders relative to low seeders at the transcriptomic level in agreement with those observed at the protein level, including a dysregulation of the adaptive immune system. The role of the immune system and neuroinflammation as a key factor in the onset and progression of neurodegenerative diseases has been widely reported^40–43^. Interestingly, a recent study using PET tracers that measure microglial activation and tau pathology in PSP patients, showed that neuroinflammation is key for progression of PSP–RS^44^ and our data would support this concept.

Taking advantage of the spatial resolution that spatial transcriptomic offers, we wanted to elucidate whether these transcriptomic changes were influenced by the presence of tau aggregates. Using a novel approach, misfolded tau protein expression was quantified and aligned with the transcriptomic data. We identified 69 genes significantly associated with tau misfolding that were enriched in negative regulation of the cell-cycle and negative regulation of transcription factor binding. Interestingly, abnormalities in cell-cycle regulation pathways have been described in neurodegenerative diseases^45,46^. Further studies on the role of these 69 genes in the pathophysiology of PSP will be needed to fully untangle the complex picture of tau aggregation and cell death.

Our multi-disciplinary approach applied in this study extends our understanding of the contribution of tau diversity to the pathology of PSP, which was previously limited to that demonstrated using conventional biochemistry and immunohistochemistry^10–14^. Our work provides strong support for the concept of molecular subtyping of PSP and, in concert with our previous work in LBD, adds to a growing literature in support of applying this concept across the neurodegenerative proteinopathies. Understanding the relationship between the seeding differences and biological activity of tau will be critical for future molecular subclassification of PSP, which, as proposed in other diseases, as cancer^47,48^ or cystic fibrosis^48^, should go beyond the conventional clinical and pathological phenotyping and consider the molecular heterogeneity of tau as well as the evaluation of other factors, such as neuroinflammation markers present in these patients. Such an approach will facilitate the development of new personalized therapeutic strategies that might need to be directed at an ensemble of differently misfolded seeding competent tau species that might also have to be combined with other immunomodulatory strategies. Furthermore, a deeper understanding of how physicochemical factors influence the aggregation of different tau polymorphs will provide support for the development of vitally needed, rapid and structure-based assays for the future diagnosis of PSP and other tauopathies.

## Material and methods

### Human tissue samples

25 (12 females, 13 males) consecutive PSP brain samples were selected from the University Health Network-Neurodegenerative Brain Collection (UHN-NBC, Toronto, Canada) based on a definite neuropathological diagnosis. Age at death, biological sex and details of a complete neuropathologic examination are provided in Supplementary Table 1.

Autopsy tissue from human brains were collected with informed consent from patients or their relatives and approval of the local institutional review board. This study was approved by the University Health Network Research Ethics Board (Nr. 20-5258).

Prior to inclusion in the study, a systematic neuropathological examination was performed following established diagnostic criteria of neurodegenerative conditions and co-pathologies^34^.

For each brain, one hemisphere was formalin fixed, prior to dissection of regions and preparation of formalin fixed paraffin embedded tissue sections required for neuropathological characterization. The contralateral hemisphere was sliced into ∼2cm thick coronal segments at the time of autopsy and immediately flash frozen and stored at −80 °C. Using a 4-mm tissue punch, microdissection of the following regions was performed: the grey and white matters of the anterior cingulate, middle frontal, primary motor, middle temporal and occipital cortices, the amygdala, hippocampus, caudate, putamen, globus pallidum, pons base, substantia nigra, subthalamic nucleus and the cerebellar white matter and dentate nucleus, as previously described^21,24^.

All the punches were stored in low protein binding tubes (Eppendorf, Hamburg, Germany), immediately flash frozen and stored at −80 °C.

Clinical data of included cases were gathered by reviewing associated patient records. Movement disorder specialists independently reviewed and extracted relevant clinical information pertaining to age at diagnosis and at death, sex, disease duration (in years), motor and non-motor symptoms, pertinent neurologic examination findings, and medications. Evolution of clinical features based on historical data was determined by documenting the temporal sequence of symptom presentation (i.e., first/presenting symptom, second symptom, third symptom). Where available, data regarding presence/absence and estimated date of onset of several motor and non-motor symptoms were collected. Age of onset of postural instability, oculomotor deficits, parkinsonism and other movement abnormalities, and cognitive impairment based on serial neurologic examination were also determined. Baseline scores on the Progressive Supranuclear Palsy Rating Scale (PSP-RS) and Montreal Cognitive Assessment (MoCA) were documented. Based on clinical features, the patients’ predominant PSP phenotype was determined using the Movement Disorder Society diagnostic criteria^6^.

### Genetic analysis

The genotypes for apolipoprotein E (*APOE*) and microtubule-associated protein tau (*MAPT*) single nucleotide polymorphisms were examined in all cases (Supplementary Table 1), using the Illumina GSA array, as previously described^16^.

### Protein extraction

For the PBS-soluble fraction, 40–50mg of frozen micro dissected tissue was thawed on wet ice and then immediately homogenized in 500μl of PBS spiked with protease (Roche) and phosphatase inhibitors (Thermo Scientific) in a gentle-MACS Octo Dissociator (Miltenyi BioTec). The homogenate was transferred to a 1.5ml low protein binding tube (Eppendorf) and centrifuged at 10,000*g* for 10 min at 4 °C, as previously described^18,21,24^. The supernatant was collected and aliquoted in 0.5-ml low protein binding tubes (Eppendorf) to avoid excessive freeze–thaw cycles. A bicinchoninic acid protein (BCA) assay (Thermo Scientific) was performed to determine the total protein concentration of all samples.

### SDS-PAGE and immunoblotting

Gel electrophoresis was performed using 4-12% NuPAGE Bis-Tris gels (Thermo Scientific). Proteins were transferred to 0.45-μm nitrocellulose membranes for 60 min at 35 V. The membranes were blocked for 60 min at room temperature in blocking buffer (5% [*w*/*v*] skim milk in 1× TBST (TBS and 0.05% [*v*/*v*] Tween-20)) or in bovine serum albumin (BSA, 5% [*w*/*v*] BSA in 1× TBST) and then incubated overnight at 4 °C with primary antibodies directed against amino acids 6-18 (Tau 12, 1:1,000 dilution, ref: 806501, Biolegend), 159–163 (Tau HT7, 1:1,000 dilution, ref: MN1000, Thermo Scientific) and 404-441 (Tau 46, 1:1,000 dilution, ref: 13-6400, Thermo Scientific) of the tau protein and against oligomeric (Tau T22, 1:2,500 dilution, ref: ABN454, SigmaAldrich) phosphorylated tau at Serine 262 (1:2,500 dilution, ref: 44-750G, Thermo Scientific), Serine 404 (1:250 dilution, ref: 44-758G, Thermo Scientific), Serine 202 and Threonine 205 (AT8, 1:1,000 dilution, ref: MN1020, Thermo Scientific), Threonine 181 (AT270, 1:10,000 dilution, ref: MN1050, Thermo Scientific) and Threonine 231 (AT180, 1:1,000 dilution, ref: MN1040, Thermo Scientific). All antibodies were diluted in blocking buffer. The membranes were washed three times with TBST and then incubated for 60 min at room temperature with horseradish peroxidase-conjugated secondary antibodies (1:3,000 dilution, ref: 172-1011 and 1:5,000 dilution, ref: 170-6515, Bio-Rad) in blocking buffer. Following another three washes with TBST, immunoblots were developed using ECL Western Blotting Detection Reagents (ref: RPN2106, Cytiva) and imaged using X-ray films. For quantification, grayscale images were scanned and imported into Fiji/ImageJ.

### 4-R Tau seeding amplification assay (4-R SAA)

SAA reactions were performed in 384-well plates with a clear bottom (Nunc) based on a previously published protocol with some modifications^35^. ThT fluorescence measurements (450±10 nm excitation and 480±10 nm emission, bottom read) were taken every 15 min for a period of 72 h. Each sample was tested in quadruplicate and the same positive and negative controls were added to each plate.

### Tau biosensor cells

Tau *in vitro* seeding assay was performed as previously described^18,25^. Briefly, the Tau RD P301S FRET Biosensor (ATCC CRL-3275) cells were cultured at 37 °C, 5% CO2 in DMEM, 10% vol/vol FBS, 0.5% vol/vol penicillin–streptomycin. Cells were plated on Ibidi clear-bottom 96-well plates at a density of 25,000 cells per well. Brain extracts (6μg of total protein from the PBS-soluble fraction) were then incubated with Lipofectamine 2000 (Invitrogen, final concentration 1% vol/vol) in opti-MEM for 10 min at room temperature before being added to the cells. Each brain region was tested in duplicate. After 48 h, cells were fixed with 4% PFA for 10 minutes and then Nucblue (Invitrogen) was added for 10 minutes at room temperature. Cells were imaged in 3 × 3 fields at 20× magnification using a Nikon ECLIPSE Ti2 confocal microscope. Using the object colocalization IF module on the HALO software (version 3.5, Indica Labs), the total number of cells (DAPI) in monolayer and tau aggregates (FITC) were quantified blindly to case identity to calculate the number of seeded aggregates per cell (FITC/DAPI).

### Cell culture and treatment

iCell Astrocytes (iAstro, Cellular Dynamic International) were thawed following company guidelines (iCell Astrocytes User’s Guide, Cellular Dynamic International). Cells were plated in complete maintenance medium (Cellular Dynamic International) on a poly-l-ornithine (ref: A004C, Sigma) and laminin (ref: L2020, Sigma) coated clear-bottom 96-well plate (Ibidi) at a density of 45,000 cells per well. iAstro cells were maintained and then treated with 6 μg of PBS soluble brain homogenates on day *in vitro* (DIV) 6 and 8. Untreated cells were used as control. On DIV 10, cells were prepared for immunocytochemistry or for cell viability assays.

### Toxicity assays

Cytotoxicity was determined by the presence of lactate dehydrogenase (LDH). Supernatant was collected at DIV 8 and 10, spun at 300 *g* for 5 minutes and stored at −80°C. LDH was measured using the Cytotoxicity Detection Kit according to the manufacturer’s instructions (ref: 11644793001, Roche, Basel, Switzerland) and absorbance (ABS) was measured at 490nm and 600nm using the SpectraMax i3 (Molecular Devices, San Jose, CA). LDH values were calculated using the equation (ABS 490nm –ABS 600nm) sample - (ABS 490nm –ABS 600nm) average media blank (background). Untreated samples were not set to 0 but were just close to the background levels of the media blanks.

Cell viability was determined at DIV 10 using the CellTiter-Blue Cell Viability Assay according to the manufacturer’s instructions (ref: 8080, Promega, Madison, WI). In brief, live cells were cultured with 20µl of reagent and 100µl of medium and incubated for 1 hour in standard cell culture conditions. Fluorescence was measured at 560/590nm (Ex/Em) using the SpectraMax i3 (Molecular Devices). Each biological sample was performed in triplicate for both assays.

### Immunofluorescence

Immunofluorescence was performed by first washing the cells in PBS and then fixing with 4% PFA for 10 minutes at room temperature. Cells were washed 3 times with PBS and incubated in 2% BSA in PBS for 1 hour and then in 0.1% TritonX100 + 1% BSA in PBS for 20 minutes, at room temperature. Samples were incubated with anti-GFAP (1:250 dilution, ref: ab7260, Abcam) in 1% BSA in PBS at 4°C overnight. The following day, cells were washed with PBS and incubated with AlexaFluor-488 donkey-anti-rabbit for 75 minutes at room temperature (1:500 dilution, ThermoFisher). Finally, cells were washed 4 times in PBS and incubated at room temperature with Nucblue (Invitrogen) for 10 minutes. Images were obtained on a Nikon ECLIPSE Ti2 confocal microscope (Nikon).

### Thermolysin and proteinase K digestions

Protease digestions were performed as previously described^24,26^, with minor modifications. A concentration of 25μg/ml of thermolysin and of 20μg/ml of proteinase K were added to 30μg of total protein from the PBS-soluble brain homogenates. Samples were incubated at 37 °C with continuous shaking (600 rpm) for 60 min. Thermolysin digestions were halted with the addition of EDTA to a final concentration of 2.5mM. Proteinase K digestions were halted with a final concentration of 4mM PMSF. Samples were resuspended in 1× LDS buffer (Life Technologies, Carlsbad, CA) and analyzed by SDS-PAGE followed by immunoblotting, as described above.

### Mass Spectrometry

Using a 4-mm tissue punch, a microdissection of the primary motor cortex from 10 PSP patients was performed. Samples were homogenized in modified RIPA buffer (2% SDS, 150mM NaCl, 50mM Tris pH8). Protein extraction was performed by mechanical disruption using 1.6mm stainless steel beads in a Bullet Blender (NextAdvance, Troy). Samples were incubated at 60 °C for 30 minutes and clarified by centrifugation. Tissue extracts were subjected to TCA precipitation as previously described^53^.Washed protein pellets were solubilized in 900 μl of urea buffer (8M urea, 150mM NaCl, 50mM Tris pH8, 1X Roche complete protease inhibitor). Protein quantitation was performed using Qubit fluorometry (Invitrogen). 50μg of lysate from each sample was reduced with 15mM dithiothreitol at 25°C for 30 minutes followed by alkylation with 151515 mM iodoacetamide at 25 °C for 45 minutes in the dark. Then, samples were digested with 2.5μg sequencing grade trypsin (Promega) at 37 °C overnight. The final digest volume was 0.5mL adjusted with 25mM ammonium bicarbonate. Then, the digest was cooled to 25 °C, acidified with formic acid and desalted using a Waters Oasis HLB solid phase extraction plate. Eluted samples were frozen and lyophilized. A pooled sample was made by mixing equal amounts of digested material from each sample. This pooled sample was used to generate a gas phase fractionation library.

#### DIA Chromatogram Library Generation

1μg of the pooled sample was analyzed by nano LC-MS/MS with a Waters M-class HPLC system interfaced to a ThermoFisher Exploris 480. Peptides were loaded on a trapping column and eluted over a 75μm analytical column at 350nL/min; both columns were packed with XSelect CSH C18 resin (Waters); the trapping column contained a 5μm particle, the analytical column contained a 2.4μm particle. The column was heated to 55°C using a column heater (Sonation). The sample was analyzed using 6 x 1.5hr gradients. Six gas-phase fractions (GPF) injections were acquired for 6 ranges: 396 to 502, 496 to 602, 596 to 702, 696 to 802, 796 to 902, and 896 to 1002. Sequentially, full scan MS data (60,000 FWHM resolution) was followed by 26 x 4 m/z precursor isolation windows, another full scan and 26 x 4 m/z windows staggered by 2 m/z; products were acquired at 30,000 FWHM resolution. The automatic gain control (AGC) target was set to 1e6 for both full MS and product ion data. The maximum ion inject time (IIT) was set to 50ms for full MS and dynamic mode for products with 9 data points required across the peak; the NCE was set to 30.

#### Sample Analysis

Samples were randomized for acquisition. 1μg per sample was analyzed by nano LC/MS with a Waters M-class HPLC system interfaced to a ThermoFisher Exploris 480. Peptides were loaded on a trapping column and eluted over a 75μm analytical column at 350nL/min; both columns were packed with XSelect CSH C18 resin (Waters); the trapping column contained a 5μm particle, the analytical column contained a 2.4μm particle. The column was heated to 55°C using a column heater (Sonation). Samples were analyzed using a 1.5hr gradients. The mass spectrometer was operated in data-independent mode. Sequentially, full scan MS data (60,000 FWHM resolution) from m/z 385-1015 was followed by 61 x 10 m/z precursor isolation windows, another full scan from m/z 385-1015 was followed by 61 x 10 m/z windows staggered by 5 m/z; products were acquired at 15,000 FWHM resolution. The maximum ion inject time (IIT) was set to 50ms for full MS and dynamic mode for products with 9 data points required across the peak; the NCE was set to 30.

#### Data processing

Data were processed with MaxQuant version 1.6.14.0 (Max Planck Institute for Biochemistry, Germany)^54^. The MaxQuant output was further processed using the Perseus software (version 1.6.14.0)^55^. Proteins with a cut-off threshold value higher than 1.33 or lower than 0.77 were considered as differential expressed proteins. All mass spectrometry data files have been deposited to the ProteomeXchange Consortium^56^ via the PRIDE partner repository. The identification of significantly dysregulated regulatory/metabolic pathways in the SN proteomic dataset was made through Metascape platform^57^ using default settings (minimum overlap: 3, minimum enrichment: 1.5, P<0.01). The functional protein association network analysis was performed using NetworkAnalyst^58^. NFT proteome and pTau interactors were obtained from *Drummond E et al*^27^.

### Histological analysis of tau pathologies

4 µm thick formalin-fixed paraffin-embedded tissue sections of neocortical regions (frontal middle gyrus, primary motor cortex, parietal inferior and superior gyrus, temporal inferior, middle, and superior gyrus, occipital cortex), basal ganglia (anterior and posterior portion), thalamus and subthalamic nucleus, hippocampus, amygdala, midbrain, pons, medulla oblongata, and cerebellum with dentate nucleus, were examined. In addition to Hematoxylin and Eosin-Luxol Fast Blue, a mouse monoclonal antibody against phospho-tau AT8 (pS202/pT205, 1:1,000 dilution, ref: MN1020, Thermo Scientific) was used for immunohistochemistry. The EnVision detection kit, Peroxidase/DAB, Mouse (Dako) was used to visualize antibody immunoreactivity. For semi-quantitative analyses of observed tau cytopathologies (neuronal, oligodendroglial, astrocytic), we used a 4-point scale: 0, absent; 1, mild; 2, moderate; and 3, severe density, as previously described^16,24^. For the figure, images were acquired using a Nikon Eclipse Ci microscope, equipped with a DS-Fi3 microscope camera and NIS-Elements imaging software (Version 1.10.00; Nikon Instruments)

### VISIUM Spatial Transcriptomics

#### Sample collection and sequencing

Primary motor cortex formalin-fixed paraffin-embedded (FFPE) tissue blocks from 4 PSP patients with DV200 > 50% were selected for sectioning. Appropriate-sized sections were placed within the frames of capture areas on the Visium Spatial Gene Expression Slide (PN-1000188, 10x Genomics) with one section in each capture area (6.5 × 6.5 mm). Tissues were deparaffinized, stained and decross-linked, followed by probe hybridization, ligation, release, and extension. Visium spatial gene expression FFPE libraries were constructed with a Visium Human Transcriptome Probe kit (PN-1000363, 10x Genomics) and Visium FFPE Reagent kit (PN-1000361, 10x Genomics) following the manufacturer’s guidance and sequenced on the Illumina NovaSeq 6000 platforms to achieve a depth of at least 25,000 mean read pairs.

#### Data Integration

Tissue quality was assessed by calculating the proportion of mitochondrial RNA for each spot, total detected gene per spot and visually inspection of tissue images. All four slices were merged, and normalized using sctransform (v0.3.3)^59^ to remove effect of library size and the proportion of mitochondrial reads. The top 2000 highly variable genes were identified using Seurat (v4.1.1)^60^, and batch effects were removed using canonical correlation analysis (cca). The integrated data was clustered using Louvain clustering with a resolution parameter of 0.8. Clusters were annotated using cell-type specific markers from publicly available human AD data^61^ and manual inspection.

#### Differential Expression

Differential expression between high- and low-seeder samples was calculated using a negative binomial mixed effect model with seeder-type and the spatial domains (Louvain clusters) as fixed effects and VISIUM slice as a random effect, as implemented in nebula (v1.4.2)^62^. Significance was assessed using 5% FDR to control for multiple testing. Genes expressed in fewer than 10% of tissue spots were excluded due to lack of power.

#### Quantification of tau load in Visium-analyzed regions of interest

For correlation of Visium gene expression data with the pathological tau load, the immediate adjacent sections (4-µm thickness) were stained against phospho-tau AT8 (pS202/pT205, 1:1,000 dilution, Thermo Scientific) using the Autostainer Link 48 (Dako). Stained sections were scanned using the Huron TissueScope LE120 (Huron) whole slide scanner and the scanned images were then collated with that of the Visium assay (10x Genomics), obtained from the Loupe Browser (10X Genomics). In the HALO software (version 3.5, Indica Labs), a 25 x 25 region of interest (ROI)-replica of the Visium assay was created using the annotation tool (each ROI at 55um in diameter and 100um distance between adjacent centers). Each ROI was annotated into individual layers (Layer 1 – 625) to obtain distinct quantification values. Using the collated image described above as a reference, the annotated template was then appropriately placed inside the grey matter at the central region of the motor cortex on the AT8-stained serial sections, placing the ROIs on top of the Visium ROIs. Tau load in each ROI was quantified using the area quantification module, reporting value as %-positive tissue area.

Differential expression in relation to tau protein expression as estimated from the matched immunohistochemical image was calculated on only the portions of two slices where tau expression was reliably estimated. Quantitative tau expression was the fixed effect and tissue slice was included as a random effect. In addition, we binned tau expression into 4 classes, which was recoded as an ordinal predictor: No tau = 0, Low tau = 1, Medium tau = 2, High tau = 3, and substituted for the quantitative estimate of tau in the model. Due to the low power (two sections only) spatial domain was not included in these comparisons.

#### Pathway Enrichments

Reactome pathways were downloaded and used for gene expression enrichment analysis (GSEA) as implemented in the fgsea (v1.20.0) package^63^. Pathways containing fewer than 10 genes, or more than 500 genes were excluded to improve power and specificity of enrichments. Genes were ranked by their estimated log2 fold-change and enrichment scores and p-values were calculated for each pathway using 50,000 permutations to assess significance. Multiple testing was controlled using a 5% FDR.

### Statistics and reproducibility

Statistical analyses were performed using GraphPad Prism (v.9) with a significance threshold of *P* = 0.05. 4R-SAA relative fluorescence responses were also analyzed and plotted using the software GraphPad Prism (v.9). The Spearman correlation coefficient *r* and the *P* value are indicated. Unsupervised hierarchal cluster analyses of the PSP cases examined were performed using SPSS Statistics (v.23).

Data collection and analyses were not performed blinded to the conditions of the experiments. For 4R-SAA analysis, all the brain regions were analyzed in quadruplicate. For the PK and the TL digestion, the same PSP brains were analyzed. For tau biosensor cells and iAstrocytes cells, all the samples were measured in duplicates.

## Acknowledgements

The authors would like to acknowledge the patients and their families for their donation. We would also like to acknowledge Dr. Michael Ford and MS Bioworks for their support with the mass spectrometry and Farzaneh Aboualizadeh from the Princess Margaret Genomic Centre and Rami Michael from 10x Genomics for their support with the spatial transcriptomics.

