## Supplementary Table, Supplementary Fig. for "4R-Tau seeding activity unravels molecular subtypes in patients with Progressive Supranuclear Palsy"

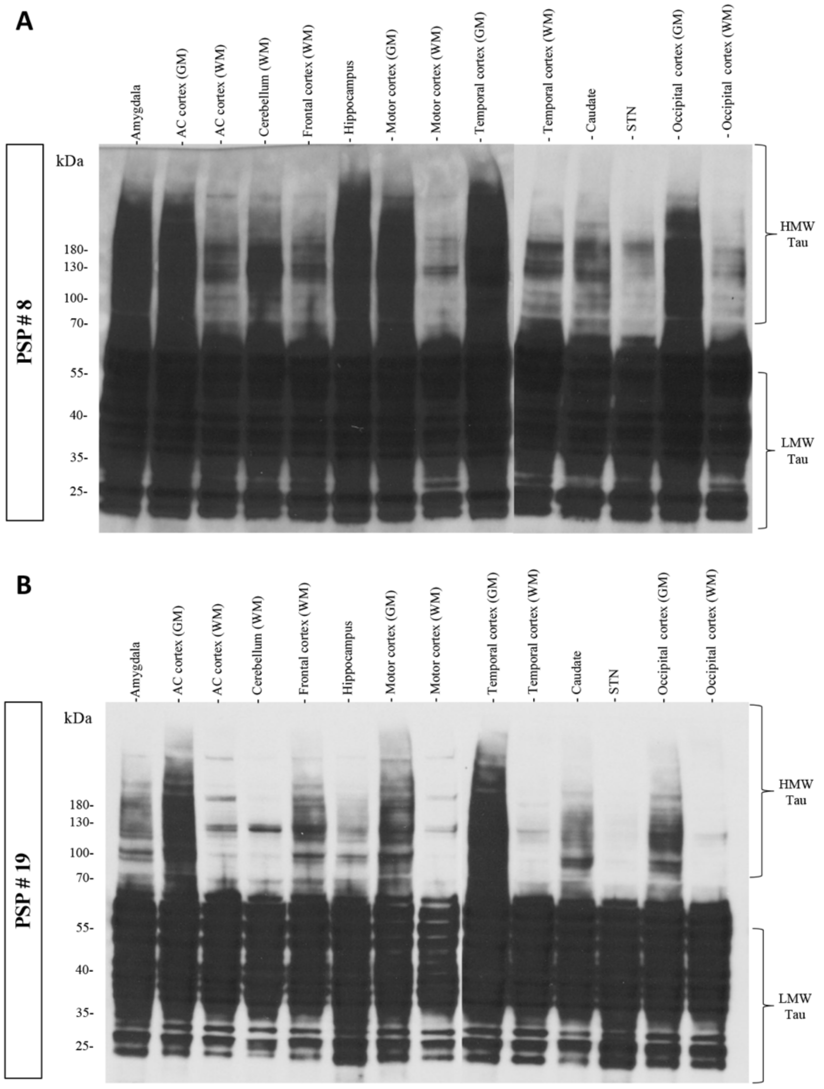


**Fig S.1 Quantification of HMW-tau reveals extensive heterogeneity between PSP patients.** ***A)*** and ***B)*** depict representative total tau immunoblots (clone HT7) showing the presence of high (> 70 kDa) and low (< 55 kDa) molecular weight tau from the motor cortex PBS-soluble fraction of two different PSP patients. HMW: high molecular weight: LMW: low molecular weight.

**A**

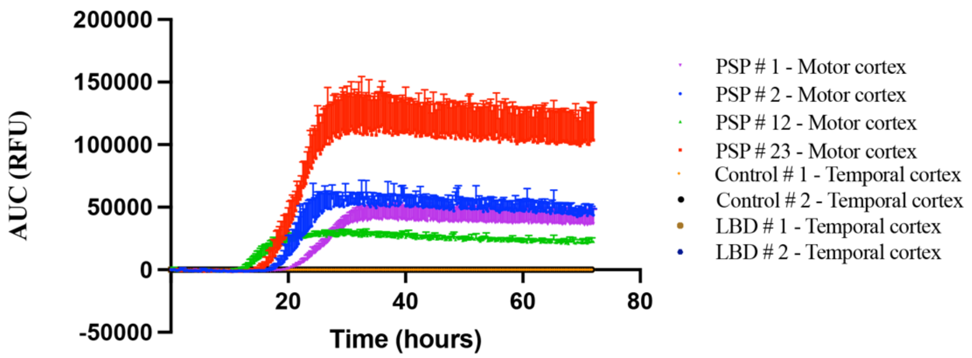


**B**

| Case | Sex | Age | NIA-AA | Cerebral Amyloid Angiopathy | Lewy Body Pathology | Lewy Body Braak Stage |
|---|---|---|---|---|---|---|
| **Control 1** | M | 74 | - | - | - | - |
| **Control 2** | F | 48 | - | - | - | - |
| **LBD 1** | M | 71 | A2B2C3 | - | Neocortical (diffuse) | Stage 5 |
| **LBD 2** | M | 79 | A3B3C3 | Aβ-positive Type 1 and 2 | Neocortical (diffuse) | Stage 5 |

**Figure S.2. 4R-Tau SAA is specific to 4R-Tau seeding. A)** Aggregation curves of 4R-Tau in the presence of motor cortex or temporal cortex homogenates from 2 healthy controls, 2 LBD cases with AD copathology and 4 PSP. Each curve represents the mean fluorescence of an individual biological sample measured in quadruplicate. **B)** Neuropathological data of the non-PSP subjects included in this assay. Lew body pathology was determined following recent Neuropathological consensus criteria for the evaluation of Lewy pathology in post-mortem brains: see: *Attems J, et al. Acta Neuropathol. 2021 Feb;141(2):159-172.* M: male; F: female; NIA-AA: National Institute on Aging and Alzheimer's Association.

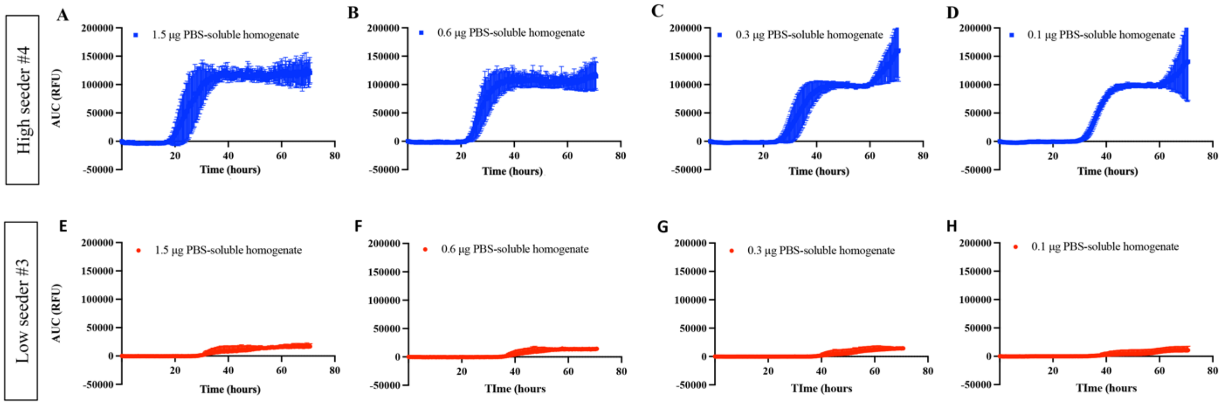


**Figure S.3: End-point dilution 4R-Tau SAA analysis of the nigral PBS-soluble fraction from two cases one classified as a high seeder and the other a low seeder.** Kinetic curves of 4R-Tau seeding activity measured by SAA of **a)** 1.5µg, **b)** 0.6µg, **c)** 0.3µg and **d)** 0.1µg of the PBS-soluble fraction from a PSP classified as high seeder and of **e)** 1.5µg, **f)** 0.6µg, **g)** 0.3µg and **h)** 0.1µg of the PBS-soluble fraction from a PSP classified as low seeder from the motor cortex. Each curve depicts the mean of the four replicates made per condition.

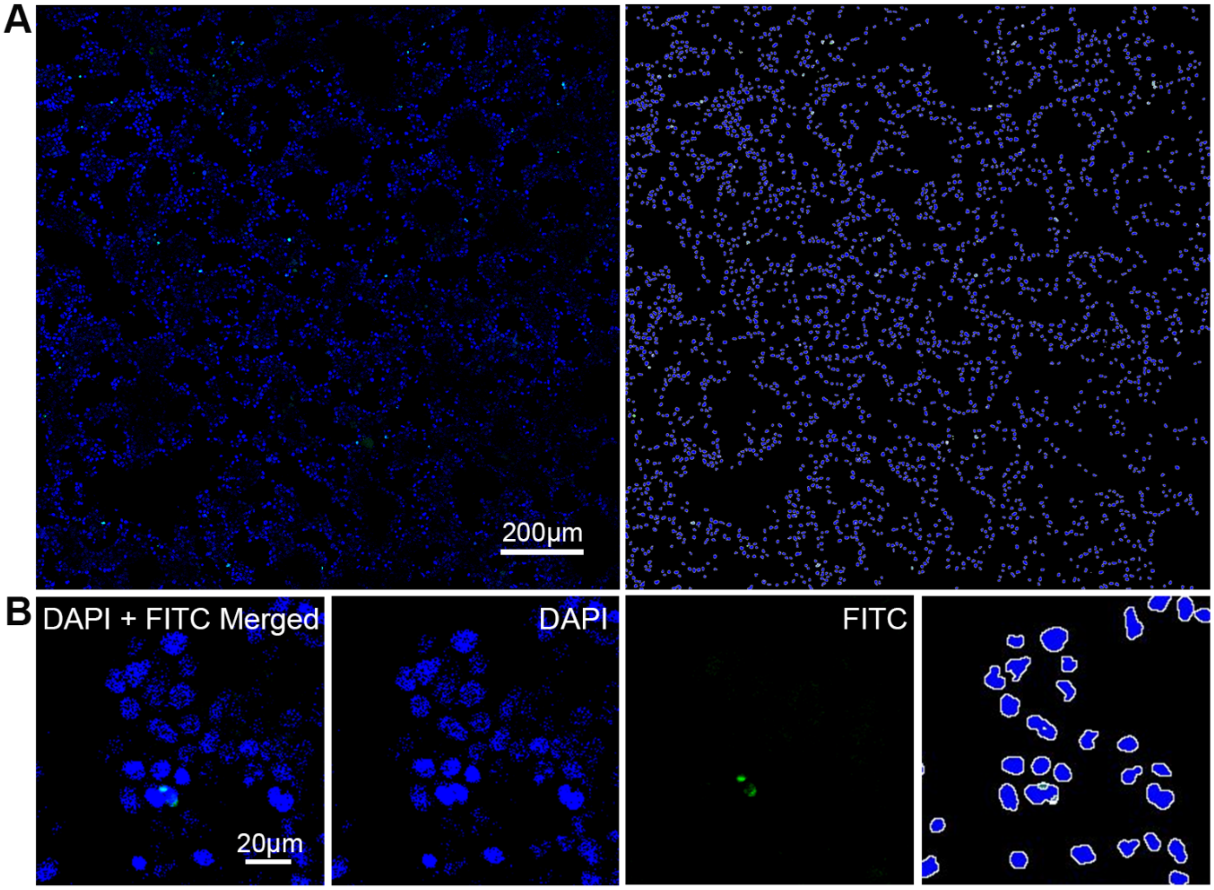


**Figure S.4 Quantification of human PSP brain-derived tau seeding using biosensor cells. A)** Cells are imaged 3x3 fields at 20x magnification for quantification of tau seeding (left). Tau aggregates fluorescence in green, which are captured using the HALO software (right; analysis mark-up image). **B)** Higher magnification image showing a tau-seeded cell. Analysis mark-up image capturing tau aggregates on far right.

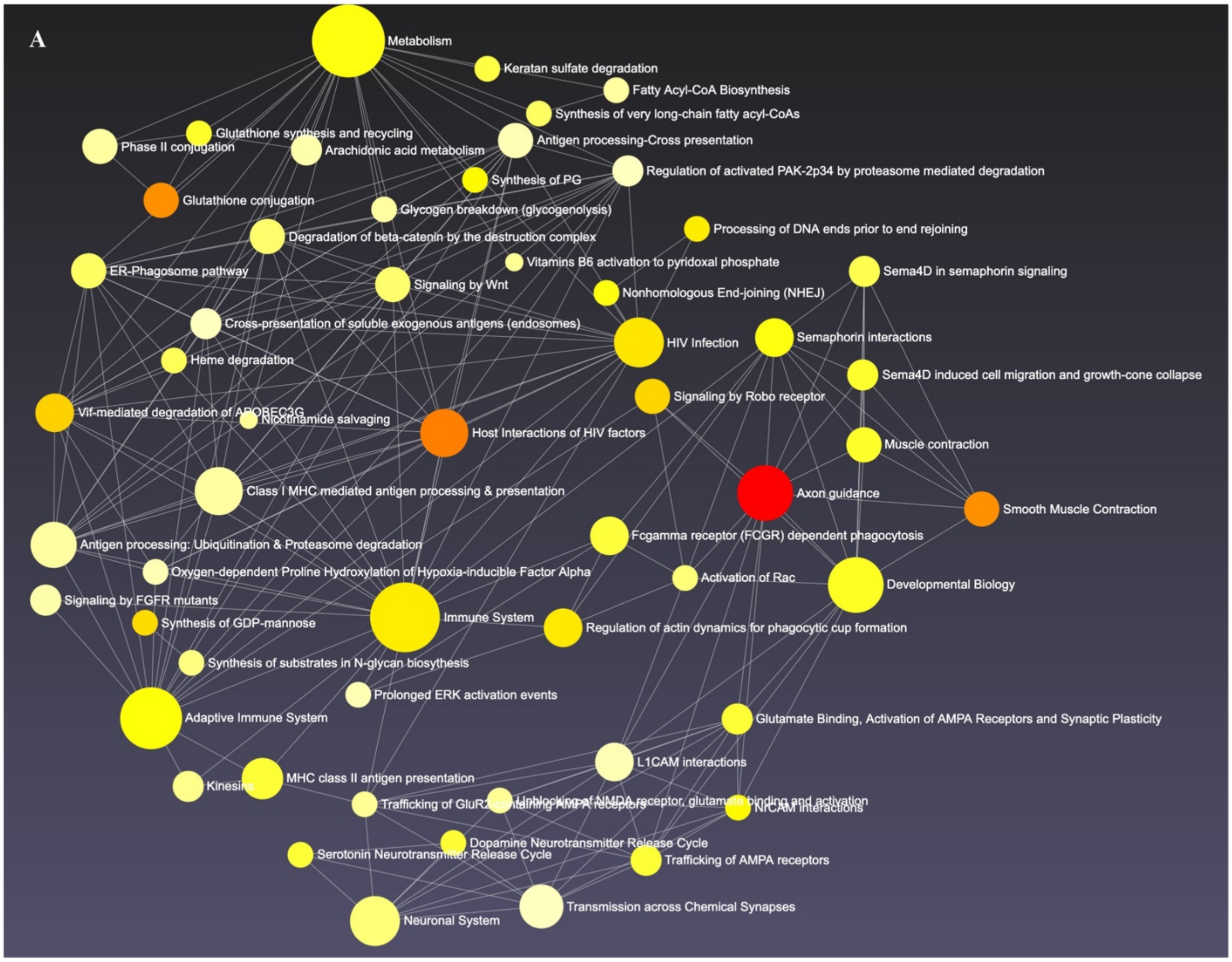


**Figure S.5 Global functional network. A)** A functional network was based on differential protein set overrepresentation generated by NetworkAnalyst and using Reactome as a pathway browser. Node size is proportional to protein sets included in each biological pathway".

| Case | Seeding category | Age | Sex | Duration of disease | PSP phenotype | PSP stage | Thal phase | NFT Braak stage | CAA: Cerebral amyloid angiopathy | LBD stage (Braak) Lewy-related pathology | TDP-43 pathology (LATE) |
|---|---|---|---|---|---|---|---|---|---|---|---|
| **PSP#01** | | 79 | M | NA | NA | 3 | Phase 3 | Stage II | Aβ-positive Type 1 | 0 | 0 |
| **PSP#02** | | 81 | M | 3 | PSP-RS | 3 | Phase 2 | Stage III | Aβ-positive Type 1 | Stage 3 | 0 |
| **PSP#03** | | 77 | F | NA | NA | 3 | Phase 4 | Stage II | No | Amygdala only | 1 |
| **PSP#04** | | 84 | F | NA | NA | 3 | Phase 4 | Stage II | Aβ-positive Type 1 | 0 | 0 |
| **PSP#05** | **HS#1** | 74 | F | 6 | PSP-RS | over 4* | Phase 2 | Stage III | No | 0 | 0 |
| **PSP#06** | | 73 | F | 9 | PSP-P | 2 | 0 | Stage II | No | 0 | 0 |
| **PSP#07** | **HS#2** | 68 | M | 4 | PSP-RS | 5 | 0 | Stage I | No | 0 | 0 |
| **PSP#08** | | 74 | F | NA | PSP-RS | 5 | 0 | Stage II | No | 0 | 0 |
| **PSP#09** | | 65 | F | NA | NA | 3 | 0 | Stage I | No | 0 | 0 |
| **PSP#10** | | 82 | F | NA | NA | 6 | Phase 3 | Stage II | Aβ-positive Type 1 | Amygdala only | 3 |
| **PSP#11** | **LS#1** | 89 | M | 10 | PSP-RS | 4 | Phase 2 | Stage III | No | 0 | 0 |
| **PSP#12** | **LS#2** | 82 | F | 9 | PSP-RS | over 4* | Phase 3 | Stage III | No | 0 | 0 |
| **PSP#13** | | 66 | F | NA | NA | 3 | 0 | Stage I | No | 0 | 0 |
| **PSP#14** | | 73 | F | NA | PSP-RS | 4 | 0 | Stage I | No | 0 | 0 |
| **PSP#15** | | 78 | M | 8 | PSP-RS | over 4* | Phase 1 | Stage II | No | Stage 1 | 1 |
| **PSP#16** | **LS#3** | 74 | M | 10 | PSP-RS | 4 | Phase 1 | Stage II | No | 0 | 1 |
| **PSP#17** | **LS#4** | 70 | F | 9 | PSP-RS | 5 | 0 | Stage I | No | 0 | 0 |
| **PSP#18** | | 90 | F | >20 | PDD | 1 | Phase 3 | Stage III | Aβ-positive Type 1 | Stage 4 | 0 |
| **PSP#19** | | 76 | M | 5 | PSP-RS | 3 | 0 | Stage II | No | 0 | 0 |
| **PSP#20** | **HS#3** | 73 | M | 3 | PSP-RS | 4 | 0 | Stage II | Aβ-positive Type 1 | 0 | 0 |
| **PSP#21** | **HS#4** | 71 | M | 6 | PSP-RS | 4 | 0 | Stage I | No | 0 | 0 |

| | | | | | | | | | | | |
|---|---|---|---|---|---|---|---|---|---|---|---|
| **PSP#22** | | 77 | M | 7 | PSP-L | 4 | Phase 1 | Stage II | No | 0 | 0 |
| **PSP#23** | **HS#5** | 72 | M | 3 | PSP-RS | 5 | 0 | Stage II | No | Stage 2 | 0 |
| **PSP#24** | | 79 | M | 7 | PSP-RS | 4 | Phase 1 | Stage III | Aβ-positive Type 1 | Stage 1 | 0 |
| **PSP#25** | **LS#5** | 69 | M | 8 | PSP-RS | 4 | Phase 1 | Stage I | No | 0 | 0 |

**Supplementary Table 1. Neuropathological and clinical data of the subjects included in this study.** *no occipital cortex was available in these cases to perfrom a complete PSP staging. HS: high seeder, LS: low seeder, NA: not available, NFT: neurofibrillary tangle, PSP: progressive supranuclear palsy.

| Case | Seeding category | ApoE | MAPT haplotype | rs564309 (TRIM11) | rs2242367 (SLC2A13 gene) | AUC 4R-Tau SAA | Tau biosensor cells |
|---|---|---|---|---|---|---|---|
| **PSP#01** | | **2 4** | H1/H2 | AA | GG | 1,801,353 | 6% |
| **PSP#02** | | **3 4** | H1/H1 | CC | AG | 2,478,246 | 5% |
| **PSP#03** | | **3 4** | H1/H1 | AC | AG | NA | NA |
| **PSP#04** | | **3 4** | H1/H1 | CC | AG | NA | NA |
| **PSP#05** | **HS#1** | **3 4** | H1/H1 | CC | AG | 3,033,167 | 21% |
| **PSP#06** | | **2 4** | H1/H1 | CC | AG | 2,232,114 | 5% |
| **PSP#07** | **HS#2** | **3 3** | H1/H1 | CC | GG | 2,762,521 | 7% |
| **PSP#08** | | **3 3** | H1/H1 | CA | GG | 2,336,660 | 8% |
| **PSP#09** | | **3 3** | H1/H1 | CC | AA | 2,500,015 | 36% |
| **PSP#10** | | **3 4** | H1/H1 | AC | GG | 2,576,900 | 18% |
| **PSP#11** | **LS#1** | **3 3** | H1/H1 | CC | GG | 1,731,695 | 3% |
| **PSP#12** | **LS#2** | **3 4** | H1/H1 | CC | GG | 1,408,088 | 3% |
| **PSP#13** | | **3 3** | H1/H1 | NA | NA | NA | NA |
| **PSP#14** | | **3 3** | H1/H1 | NA | NA | NA | NA |
| **PSP#15** | | **3 3** | H1/H1 | AC | AA | 1,979,323 | 2% |
| **PSP#16** | **LS#3** | **3 4** | H1/H1 | CC | AG | 888,331 | 2% |

| | | | | | | | |
|---|---|---|---|---|---|---|---|
| **PSP#17** | **LS#4** | **3 3** | H1/H1 | AC | GG | 1,795,747 | 0% |
| **PSP#18** | | **3 4** | H1/H1 | AC | AG | 285,861 | 0% |
| **PSP#19** | | **2 3** | H1/H1 | AC | GG | 1,817,046 | 7% |
| **PSP#20** | **HS#3** | **3 3** | H1/H1 | CC | AG | 2,950,152 | 3% |
| **PSP#21** | **HS#4** | **3 3** | H1/H1 | CC | AG | 3,192,424 | 24% |
| **PSP#22** | | **3 3** | H1/H1 | CC | GG | 2,701,916 | 11% |
| **PSP#23** | **HS#5** | **3 3** | H1/H1 | CC | GG | 5,381,318 | 14% |
| **PSP#24** | | **3 3** | H1/H1 | CC | AG | 2,644,363 | 16% |
| **PSP#25** | **LS#5** | **3 3** | H1/H1 | CC | AG | 1,518,539 | 9% |

**Supplementary Table 2. Genetic and Tau seeding data of the subjects included in this study. AUC: area under the curve,** HS: high seeder, LS: low seeder, NA: not available, PSP: progressive supranuclear palsy, SAA: seeding amplification assay.

| Gene | p_value | FDR |
|---|---|---|
| MT1H | 1.29E-10 | 0.00000157 |
| RAB7B | 0.00000107 | 0.001620744 |
| FOXJ1 | 0.000000389 | 0.000786703 |
| SLFN11 | 0.000255003 | 0.045518022 |
| SLC24A4 | 0.0000094 | 0.007127445 |
| SLC38A11 | 0.0000151 | 0.008709745 |
| PROX1 | 0.000000718 | 0.001244363 |
| CASP7 | 0.000152907 | 0.031657343 |
| SMTN | 9.66E-09 | 0.0000586 |
| GSDMB | 0.000031 | 0.012129764 |
| LRATD1 | 0.000153879 | 0.031657343 |
| TRIO | 0.0000246 | 0.011071742 |
| DDIT3 | 0.000216906 | 0.040504737 |
| SH3GL1 | 0.0000285 | 0.011518045 |
| ANGEL2 | 0.000118574 | 0.028220501 |
| TENT4B | 0.0000117 | 0.007136285 |
| IRF2BP2 | 0.0000336 | 0.01236781 |
| SLC30A4 | 0.000135656 | 0.030290038 |
| RTL5 | 0.000152622 | 0.031657343 |
| BMP7 | 0.000064 | 0.02004766 |
| TMEM108 | 0.0000881 | 0.022760385 |
| ZNF141 | 0.0000716 | 0.020214169 |
| VEZF1 | 0.0000419 | 0.014529007 |
| ID1 | 0.0000871 | 0.022760385 |
| PAPOLG | 0.0000707 | 0.020214169 |
| CYP20A1 | 0.0000195 | 0.010302584 |
| PACSIN3 | 0.000270012 | 0.047498569 |
| CD82 | 0.000023 | 0.010755358 |
| GNB1L | 0.00000386 | 0.004261636 |
| KDM6B | 0.0000796 | 0.021954219 |
| YBEY | 0.0000713 | 0.020214169 |
| SNX11 | 0.00000494 | 0.004996297 |
| TOP3B | 0.0000476 | 0.016033765 |
| GIMAP7 | 0.000111159 | 0.026984847 |
| PHTF2 | 0.00000864 | 0.006987762 |

| Gene | p_value | FDR |
|---|---|---|
| SMC4 | 0.000194596 | 0.037492125 |
| KATNAL2 | 0.00000999 | 0.007131427 |
| HIC2 | 0.00014589 | 0.031621635 |
| CEP57 | 0.00000206 | 0.002781425 |
| TNS3 | 0.0000501 | 0.016440119 |
| FILIP1L | 0.0000667 | 0.020214169 |
| ANXA2 | 0.000240681 | 0.043602797 |
| DTWD1 | 0.000103237 | 0.025573384 |
| HIST1H4J | 0.0000258 | 0.011179324 |
| GLIS2 | 0.0000403 | 0.014391522 |
| MMP15 | 0.0000227 | 0.010755358 |
| PEX11A | 2.87E-08 | 0.000101487 |
| LNX2 | 0.000235649 | 0.043338041 |
| SNX18 | 0.0000281 | 0.011518045 |
| MKS1 | 0.000124955 | 0.028617084 |
| ZNF207 | 0.00000621 | 0.005801648 |
| MOG | 0.000192182 | 0.037492125 |
| NPPC | 0.0000836 | 0.022558736 |
| BAMBI | 0.0000118 | 0.007136285 |
| GPAT3 | 0.00000317 | 0.003843561 |
| IPMK | 0.000158555 | 0.0320756 |
| TNNI3K | 0.000022 | 0.010755358 |
| ENKUR | 0.00000857 | 0.006987762 |
| ZNF805 | 3.34E-08 | 0.000101487 |
| PCDHB9 | 0.000199956 | 0.037922827 |
| CENPL | 0.000137251 | 0.030290038 |
| ACADL | 0.0000644 | 0.02004766 |
| SFT2D2 | 0.000000205 | 0.000497062 |
| TGFB1 | 0.000032 | 0.012138391 |
| MRC2 | 0.0000114 | 0.007136285 |
| HCK | 0.000168201 | 0.033469265 |
| ADAMTSL1 | 0.000123525 | 0.028617084 |
| RCAN3 | 0.0000918 | 0.02322037 |
| DEF6 | 0.0000189 | 0.010302584 |

**Supplementary Table 3.** List of genes significantly associated with tau misfolding.
